# Energetic misfires: Hybridization drives transgressive expression in metabolic pathways in thermally divergent Icelandic stickleback

**DOI:** 10.64898/2026.04.17.719192

**Authors:** Matthew K Brachmann, Bethany Smith, Bjarni Kristjánsson, Colin Selman, Kevin Parsons

## Abstract

Climate change is causing rapid changes to freshwater environments, driving selection for phenotypes that can cope with altered thermal conditions, while also changing developmental environments. This may promote local adaptation, migration to new habitats, and/or phenotypic plasticity. Climate change may also increase hybridization rates between locally adapted phenotypes, as populations migrate and spatially mix in new ways. Consequently, these conditions may facilitate the production of variation, including both adaptive and maladaptive outcomes. To examine this, we leveraged threespine stickleback (*Gasterosteus aculeatus*) that have locally adapted to either geothermally warmed or ambient environments in Iceland. We tested the effects of hybridization between thermal ecotypes on gene expression using a common garden experiment, where pure-strain ecotypes and their hybrids were reared under 12°C and 18°C. We performed RNA-seq on brain and liver to assess 1) ecotype divergence, 2) plasticity, and 3) the effects of hybridization and inheritance patterns. We identified a low degree of expression divergence between locally adapted ecotypes, despite a high degree of plasticity across rearing environments. Hybrid ecotypes were highly divergent from both geothermal and ambient ecotypes and exhibited transgressive expression under both rearing temperatures. Transgressive expression disrupted gene networks extensively, with broader effects at 18°C than at 12°C, primarily associated with metabolism and mitochondrial function. Hybridization between locally adapted thermal ecotypes appears to largely disrupt genes associated with energy balance and metabolic function. While demonstrating mechanisms underlying the rapid evolution of reproductive isolation, these findings also provide insights for how populations may cope or fail within a warming world.

## Introduction

Climate change presents an ever-worsening threat to biodiversity (IPCC, 2021) as continued warming pushes species to their thermal limits (Nunez et al., 2019). Ectotherms are expected to be particularly vulnerable due to thermoregulation being mediated by the environment (Fry, 1967; Paaijmans et al., 2013), and where increasing temperatures destabilize thermoregulatory capacity (Haesemeyer, 2020; Duffy et al., 2022). Climate change will require ectotherms to either migrate to more favorable habitats or rapidly adapt to warming environments (Parmesan & Yohe, 2003; Deutsch et al., 2008; Chen et al., 2011). In general, species experiencing the greatest amount of warming have moved larger distances with latitudinal shifts tracking temperature changes over time (Chen et al., 2011). Climate change has already been implicated as a cause for adaptive phenotypic shifts across fishes (Crozier & Hutchings, 2014). However, the contribution of phenotypic plasticity underlying such changes is unclear with a heritable basis typically being untested (Crozier & Hutchings, 2014). Thus, there is a pressing need to understand how climate change can induce adaptation through plasticity, genetic evolution, and exactly how these factors interact. Determining the underlying mechanisms is an area of growing interest as it may inform conservation strategies (Reed et al., 2011; Campbell et al., 2017; Greenspoon & Spencer, 2021; Parsons, 2021; She et al., 2023; Sheldon et al., 2025; Wighard et al., 2025).

The capacity for adaptation to climate change is dependent on the amount of phenotypic variation generated through genomic and developmental processes (Pigliucci, 2008; Campbell et al., 2017; Riederer et al., 2022). The production of adaptive phenotypic variation through development involves a combination of environmental, genomic, and regulatory changes (Hansen, 2006; Riederer et al., 2022). The inheritance of adaptive phenotypic variation can be both additive and non-additive (Falconer & Mackay, 1996), with phenotypic expression being further modified by cis- vs trans-regulatory elements (Wittkopp et al., 2004). While cis-regulatory elements act on specific alleles, trans-regulatory elements influence both alleles equally. Allele specific expression allows for the disentanglement of the inheritance of regulatory variation and differentiate between cis- and trans-regulatory effects (Fan et al., 2020; Pierre et al., 2022). Transgressive expression, due to non-additive inheritance, involves gene expression in hybrids that is greater or less than parental phenotypes (Go & Civetta, 2020; Wang et al., 2022; Yazdi et al., 2022) and can arise when cis- and trans-regulatory elements are decoupled, revealing cryptic regulatory variation (Lemmon & Juenger, 2017; Jacobs et al., 2024). The relative role of cis- vs trans-regulatory elements, and their interaction with both additive and non-additive inheritance mechanisms, in response to changing environments has been debated (Crispo et la., 2008; Lemos et al., 2008; Osada et al., 2017; McGirr & Martin, 2021).

Hybridization between locally adapted species or ecotypes may promote genomic and phenotypic variation within novel environments (Abbott et al., 2013; Brice et al., 2021; Jacobs et al., 2024). Secondary contact and hybridization may occur between locally adapted populations across both small and large scales (Chunco, 2014). Hybrid offspring at zones of secondary contact are unlikely to have phenotypes that match either parental phenotype (Gow et al., 2007; Rundle & Nosil, 2005) and these hybrid phenotypes may be selected against (Servedio, 2004; Lemmon & Kirkpatrick, 2006; Lenormand, 2012; Butlin & Smadja, 2018). Hybrids are generally predicted to be maladaptive in either parental environment due to a disruption of adaptive gene by environment interactions (Carroll et al., 2003; Cenzer, 2017) and can reinforce reproductive isolation (Lemmon & Juenger, 2017). In a warming world, populations that are locally adapted to different thermal conditions may come into contact at secondary contact zones as they migrate (Chunco, 2014). Therefore, addressing how parental and hybrid offspring of thermally adapted populations respond to temperature variation could provide new insights into understanding how genomic and regulatory mechanisms are impacted by climate change.

Icelandic threespine stickleback (*Gasterosteus aculeatus*) provide a powerful system to untangle the effects of hybridization on the molecular architecture of adaptation to warming environments. While understanding adaptation to warming environments often involves comparing populations from widely varying latitudes or altitudes as a proxy, this confounds temperature variation with other environmental factors (Daco et al., 2021; Pilakouta et al., 2023). Populations of three-spined stickleback (*Gasterosteus aculeatus*) avoid such confounds by inhabiting geothermal and ambient habitats in close geographic proximity across Iceland (Millet et al., 2013; Pilakouta et al., 2020; Pilakouta, et al., 2023). Geothermal and ambient habitats exhibit a strong thermal gradient that typically differ by ∼10°C throughout the year, but with otherwise similar environments (Pilakouta et al., 2020). Stickleback living across geothermal-ambient gradients demonstrate heritable adaptive divergence in as little as 70 years, including large fitness tradeoffs (Pilakouta et al., 2023; B. Smith, 2023; B. A. Smith et al., 2024).

To understand the genomic and developmental mechanisms underlying phenotypic divergence between thermal ecotypes we tested the influence of plasticity and hybridization on gene expression. We tested three hypotheses: 1) Geothermal and ambient stickleback ecotypes will show divergent and plastic gene expression across temperature environments, reflecting local adaptation. 2) Hybrid expression will differ from pure-strain ecotypes due to non-additive, transgressive expression. 3) Hybridization between locally adapted ecotypes will disrupt co-adapted cis-trans regulatory networks, leading to misexpression of specific alleles of locally adapted genes and maladaptive phenotypes.

## Methods

### Creation of F1 hybrid and pure strain families

Sexually mature sticklebacks were captured using un-baited minnow traps (mesh size, 6.4mm) from an allopatric geothermal-ambient habitat pair in Northern Iceland, near Sauðárkrókur (geothermal habitat: 65.732260°, −19.618937 °, ambient habitat 65.732191 °, −19.618574 °) in 2017 and brought to the University of Glasgow. Ambient and geothermal summer temperatures were simulated in the aquaria using 12°C (±1°C) and 18°C (±1°C), respectively. Following two months of a simulated winter light cycle (4hrs light and 20 hours darkness) fish were primed for reproduction with nutrient rich food (bloodworms, mysis) and an extended light period (20-23hrs light and 4-1 hour darkness). *In vitro* fertilization was performed (Barber & Arnott, 2000) to create four ambient families, four geothermal families, and two hybrid families using two geothermal females and two ambient males (Figure 1). All egg clutches were left to fertilize for fifteen minutes before being split evenly to 12°C (±1°C) and 18°C (±1°C).

**Figure 1.**
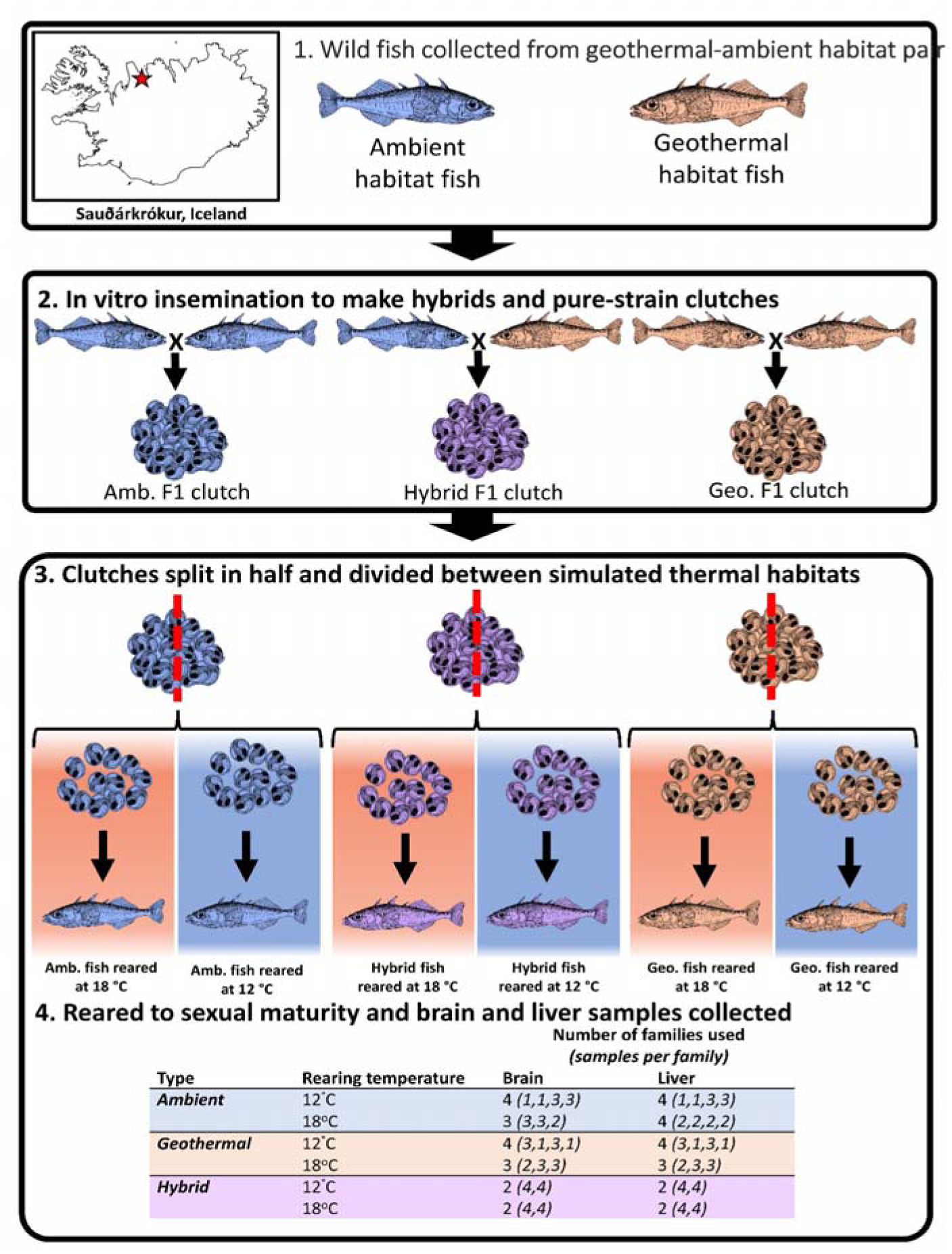
Methodological approach for the creation of experimental hybrid and pure-strained crosses of geothermal and ambient Icelandic threespine stickleback. 1. Wild geothermal and ambient stickleback were collected from allopatric Saudárkrókur (SKR) populations in Iceland. 2. Hybrid and pure-strained geothermal and ambient crosses were made via invitro insemination. Eggs were striped from females and fertilized with sperm from the corresponding pure-strained male. Hybrid crosses were made by crossing ambient males with geothermal females. 3. Hybrid and pure-strained crosses were divided in half and raised under 12°C and 18°C rearing conditions until maturity.

Fertilized clutches were kept in mesh baskets submerged within larger basins of water fed by a recirculated continuous water drip system. Water was treated with methylene blue (2.5 µg/ml) until hatching to reduce the risk of fungal infection (Ansari et al., 2016). Containers were inspected daily, and dead or infected eggs were removed immediately. After hatching, larvae and juveniles were fed live food (newly hatched brine shrimp (*Artemia* salina), and microworms (*Panagrellus redivivus*) as well as size-appropriate ZM powdered food (ZM100 and ZM200) (ZMsystems, Twyford, UK). Upon reaching approximately 2cm, juveniles were moved, in their family groups, to 10L tanks at standardized densities of 15-20 individuals. From 3cm to maturation fish were fed trout pellets (Microstart, EWOS Ltd, Surrey, UK). The photoperiod of the aquarium was kept at a 12:12 light cycle.

### Tissue collection, RNA extraction and sequencing

Brain and liver samples were collected from stickleback for each treatment condition. Sampled fish were one year old (±1 month), and that provide a size match across treatments. Threespine stickleback were euthanized with an overdose of benzocaine following the Home Office guidelines on Animal use. Individuals were sexed by the identification of testes and ovaries. Brain and liver samples were collected from 8 individuals per ecotype for each temperature treatment with sampling from 3-4 separate families for the geothermal and ambient ecotypes and 2 families for the hybrids, resulting in 48 total samples (Figure 1). Each tissue was cut into 0.5cm^3^ pieces and stored in RNAlater (Invitrogen AM7020) at room temperature overnight before freezing at −80°C.

To obtain RNA, extractions were performed using a Qiagen RNeasy standard kit (Qiagen Cat ID: 74104) for liver tissue (with 50% ethanol as recommended for fatty tissue such as liver) and the Qiagen RNeasy lipid tissue kit (Qiagen Cat ID: 74804) for brain tissue. RNA was extracted from ∼20mg of tissue and purity was determined using a NanoDrop spectrophotometer (Thermo Fisher Scientific, Massachusetts, US), where all samples had 260/230 or 260/280 ratios > 1.8. Concentration was determined using a RNA Broad Range kit (Thermo Fisher Scientific, Massachusetts, US) on a Qubit 4 Fluorometer using a Qubit RNA Broad Range kit (Thermo Fisher Scientific, Massachusetts, US). RNA integrity was assessed for all samples, using an Agilent 4200 Tapestation (Agilent Technologies, California, US), to calculate RNA integrity number (RIN values). RIN values are a numerical score ranging from 1-10 where a higher value indicates a higher degree of RNA integrity (Sheng et al., 2017) and all samples has a RIN value >7.

The preparation of cDNA libraries and whole transcriptome sequencing were performed by the Center for Genomic Research (CGR) a the University of Liverpool using the NEBNext® Poly(A) mRNA Magnetic Isolation Module (NEB E7490) and NEBNext® Ultra RNA Library Prep Kit for Illumina® (E7530). Sequencing was performed on the Novaseq 6000 platform (Illumina, California, US) aiming for an average read length of 150 base pairs and 20M sequenced reads per sample.

### RNA-seq data analysis

#### Initial data processing and quality control

Read quality was investigated using *FastQC* (Andrews, 2010) and *MultiQC* (Ewels et al., 2016). Fastq files were trimmed for the presence of Illumina adapter sequences using *Cutadapt* version 1.2.1 (Martin, 2011) and further trimmed using *Sickle* version 1.200 (Joshi & Fass, 2011) before alignment to version five of the three-spined stickleback reference genome (Genbank GCA_016920845.1) (Peichel et al., 2020) using the *Hisat2* aligner version 2.2.0 (Kim et al., 2019). The resulting SAM files were then sorted and converted to BAM format using *Samtools sort* and indexed using *Samtools index* (Danecek et al., 2021). Read counts were obtained using *Htsqcount2* (Anders et al., 2015).

#### Differential expression between ecotypes and temperatures

Data analysis was performed in R (v4.5.1) (R Core Team, 2022). The packages *EdgeR* (Robinson et al., 2009) and *LIMMA* (Ritchie et al., 2015) were used in parallel to provide higher confidence through corresponding results. The read count data was filtered for low-expressed genes using the *filterbyExpr* function in the *EdgeR* package. Genes were filtered out if they had fewer than ten counts per million (CPM) in more than 8 samples for each tissue type (Y. Chen et al., 2016). Read counts were normalized by converting observed library sizes into effective library sizes using scaling factors calculated by *calcNormFactors* using the TMM method (Robinson & Oshlack, 2010).

To address our first hypothesis, we tested for differences in gene expression between ecotypes, hybrids, rearing temperatures, and their interaction using a single contrast matrix. For all analyses the false discovery rate (FDR) was controlled using the Benjamini-Hochberg method (Benjamini & Hochberg, 1995). Genes were considered differentially expressed when the resulting adjusted p value was below 0.05. In our contrast matrix, we tested for divergent expression by subtracting expression values for geothermal fish at a given rearing temperature from ambient expression at the same temperature, resulting in two contrasts (“*ambient at 12°C – geothermal at 12°C”* and “*ambient at 18°C – geothermal at 18°C”*). To examine plasticity we compared expression at 18°C versus 12°C within each fish type (geothermal, ambient, and hybrid). To address our second hypothesis, we then tested for mismatches between hybrids and pure ecotypes by subtracting hybrid expression at each temperature from a given pure ecotype expression at the same temperature, resulting in four contrasts (“*ambient at 12°C – hybrid at 12°C”, “ambient at 18°C – hybrid at 18°C”, “geothermal at 18°C – hybrid at 18°C”,* and *“geothermal at 12°C – hybrid at 12°C”*). In *EdgeR* a negative binomial generalized linear model was fitted to each gene. This analysis assumed that the variance of gene counts was dependent on the negative binomial dispersion and the quasi-likelihood dispersion (Ren & Kuan, 2020). Here, the negative binomial dispersion represented the variability of the biological system, while the quasi-likelihood dispersion represented gene-specific variability greater or lower than the overall level, capturing both biological and technical sources of variability (Lun et al., 2016). The mean-dispersion across all genes was used to estimate the negative binomial dispersion using the *estimateDisp* function (Lun et al., 2016; Ren & Kuan, 2020). An empirical Bayes approach was then used to estimate the quasi-likelihood dispersion and a generalized linear model accommodating the design model matrix was fitted using *glmQLFit* (Lun et al., 2016; Ren & Kuan, 2020). The top-ranked genes for each contrast were then extracted using *TopTags*.

A complementary analysis using *LIMMA* was then performed to test differences in gene expression within our four contrast matrices. Normalized read counts were prepared for linear modelling in *LIMMA* by performing a voom transformation, which incorporated the mean-variance relationship of the log-counts into the precision weights for each observation (Law et al., 2014, 2018). Next, a linear model was fitted to each gene using *lmFit* (Law et al., 2018). A corollary test was then done using *eBayes* to perform a moderated F-statistic test and rank genes by evidence of DE for each contrast (Law et al., 2018). The top-ranked genes for each contrast were then extracted using *topTable*.

We then compared the results from *EdgeR* and *LIMMA* and selected genes that showed differential expression in both analyses. For both approaches the fitted models used contrast matrices designed to address our questions around divergence, plasticity, and their interaction. These contrast matrices described the linear combinations of parameters used to calculate the differences between groups of interest (Law et al., 2020). We then tested for differences between fish types and rearing temperatures using a principal component analysis (PCA) on all differentially expressed genes for brain and liver tissues using the *prcomp* function.

### Hybrid gene expression and inheritance mode categorization

To further address our second hypothesis, we assessed inheritance mode as it was necessary to categorize patterns of gene expression. Thus, in each rearing temperature hybrid expression was compared to pure strain fish. Genes with DE in at least one of the previous hybrid to pure-strain contrasts were selected for comparison, and inheritance modes were classified separately for each rearing temperature. Specifically, genes were categorized based on the log fold changes (LFC) of DE between hybrids and each pure-strain type with a threshold of 0.32 (1.25-fold difference, shown to be a useable cut-off by (Wang et al., 2022; Yazdi et al., 2022)). DE genes were categorized as either additive, dominant, or transgressive on the basis of their respective LFC threshold (Figure 2) Dominant gene expression was assessed for each of the ecotypes where the hybrids had a LFC between −0.32 and 0.32, we could then assess dominance due to either the geothermal or ambient ecotype. Additive gene expression was classified as being >0.32 LFC in one ecotype and <0.32 in the other ecotype. Lastly, transgressive gene expression was classified as the LFC being > or < 0.32 in both ecotypes.

**Figure 2.**
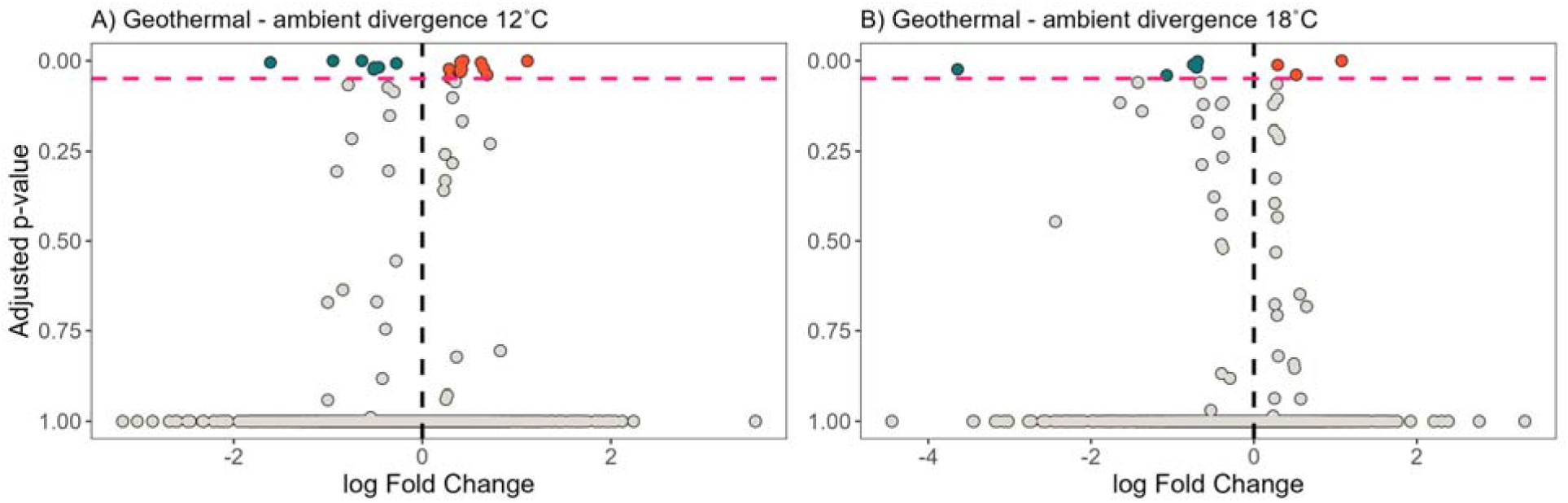
Volcano plots showing differentially expressed genes between the geothermal and ambient pure-strained ecotypes under 12°C (A) and 18°C (B). The red dashed line represents the p-value cut-off of 0.05. Genes in blue are down-regulated and genes in orange are up-regulated.

### Testing for divergence and disruption in gene co-expression

To assess our third hypothesis, we assessed the effects of hybridization on gene co-expression networks linked up developmental pathways (Oldham et al., 2006). Changes in gene networks between ecotypes can be used to discover key drivers of evolutionary divergence (Oldham et al., 2006), while hybrids can display disrupted co-expression networks that may alter phenotypic development and decrease fitness (Filteau et al., 2013). Thus, gene networks associated with divergence and plasticity were identified using the *GeneCoEx* workflow in R (Li et al., 2023) whereby differentially expressed genes for each contrast and tissue type were used to produce a correlation matrix and detect co-expression modules. First, significant DEGs from each contrast were filtered to remove genes with invariable expression profiles. A correlation matrix was then made for all genes and correlations greater than 0.7 were retained and relationships between highly correlated genes were calculated. For each contrast, modules of gene expression were calculated using the *cluster_leiden* and *optimize_resolution* functions. The *cluster_leiden* function performs Leiden clustering using the Leiden algorithm (Traag et al., 2019) to find modularity in gene expression. To control false discovery and spurious results, each module from the *optimize_resolution* function had to contain 5 or more genes with gene expression being calculated for each module. The gene network was calculated using the *graph_from_data_frame* function which uses the edges and nodes calculated with correlation coefficients greater than 0.7. Gene networks for each contrast were then imported into *Cytoscape* (Ono et al., 2025) for visualization and gene ontology analysis. Gene ontology (GO) analysis was then performed for each gene network using *Enrichment Map* within *Cytoscape* (Merico et al., 2010).

### Comparing ecotype regulatory mechanisms through allele specific expression analysis

To further address our third hypothesis, we assessed divergence in regulatory mechanisms through investigation of allele-specific expression (ASE), defined as the imbalanced expression of maternal or paternal alleles of a gene in a diploid individual. ASE can influence evolution by enabling natural selection to act on individual gene copies, allowing organisms to mask deleterious mutations or adaptively overexpress beneficial ones (Wittkopp et al., 2004, 2008; Wittkopp & Kalay, 2012). Thus, ASE can act as a regulatory mechanism that creates phenotypic variation, driving rapid adaptation to environmental changes and influence complex traits through tissue-specific and parent-of-origin effects. The premise for the identification of ASE is that one copy of each pure-strain chromosome should be present in each cell of an F1 hybrid, meaning that while the trans-regulatory environment for each allele is the same, the cis-regulatory environment may differ. Differences in cis-regulatory elements can therefore result in the differential expression of each allele in hybrids.

To assess the presence of allele specific expression, aligned Bam files were prepared for variant calling using the *GATK* best practices RNAseq short variant discovery workflow (Brouard et al., 2019; Caetano-Anolles, 2023b). Post-alignment quality was checked using *Picard ValidateSamFile* while mate-pair information was verified and fixed, and duplicate reads tagged using *Picard’s FixMateInformation* and *MarkDuplicates* (Institute, 2023). *GATK’s SplitNCigarReads* was used to reformat alignments spanning introns for compatibility with *HaplotypeCaller.* An initial round of variant calling was performed in order to bootstrap a set of known variants, which were filtered (*GATK: VariantFiltration, SelectVariants*) using hard filters (Quality by Depth (QD) <2.0, Fisher Strand (FS) > 60, RMS Mapping Quality (MQ) < 40, Strand Odds Ratio (SOR) >4, as recommended by *GATK* best practices (Caetano-Anolles, 2023a)), for input into *GATK’s Base Quality Score Recalibration*. Base quality recalibration was performed using machine learning applied to each sample to detect and correct patterns of systematic errors in the base quality scores using *BaseRecalibrator*. A second round of variant calling using *HaplotypeCaller* provided variants, which were also filtered (*VariantFiltration, SelectVariants*) using hard filters (QD <2.0, FS > 60, MQ < 40, SOR >4) and SNPs were called for all fish (geothermal, ambient, and hybrid). Pure-strain SNPs were filtered for homozygosity across each ecotype to extract those unique to each.

SNPs identified in hybrid fish were filtered to include only those that were heterozygous to target those that may be habitat divergent. Allele specific reads were then output from the VCF files using GATK’s *ASEReadCounter*. SNPs were mapped to genes using *SNPeff* and filtered for a total read count of >20 with a minimum read count for the reference and alternatives of >5. Allele specific expression was then assessed per SNP using Allele-Specific Expression analysis in a Population (ASEP) (Fan et al., 2020). ASEP is a method for gene-level detection of allele-specific expression at the population level without individual genomic data, using a pseudo-phasing procedure that uses allele-specific read counts to infer haplotype phases through a majority voting procedure based on allele specific read counts. A generalized linear mixed-effects model is then used to test for allele-specific expression. Because differing levels of allele-specific expression between rearing temperatures could indicate environmental effects on expression regulation ASEP analysis was performed separately for hybrids reared at 12°C and 18°C. To assess biological relevance SNPs identified by ASEP analysis were cross-referenced to SNPs present in pure-strain fish that were unique to either ecotype.

Genes were annotated using Ensembl three-spine stickleback gene stable IDs. To conduct ontology analysis Ensembl Stickleback gene stable IDs were converted to Ensembl Zebrafish gene stable IDs using *biomaRt* (Durinck et al., 2009; Kinsella et al., 2011). The Metascape package (Zhou et al., 2019) was then used on converted data to interpret the list of DE genes produced from each analysis (minimum overlap 3, p value cut-off 0.01 and minimum enrichment 1.5). Gene ontology (GO) enrichment analysis relies on an accurate assessment of whether a particular gene ontology is overrepresented in a dataset. Therefore, in order to avoid sampling bias, background gene sets of the expressed genes only (non-zero total read), rather than the whole zebrafish genome, were used for each tissue (Timmons et al., 2015). As the recommended maximum for Metascape gene ontology analysis is 3000 genes, gene lists that exceeded this threshold were filtered to remove genes with an absolute LFC lower than 0.5. This analysis was performed only on modules found to be significantly divergent, disrupted or plastic. The interpretation of module gene ontology was further simplified by selecting the top-level gene ontology terms containing the most significant lower level GO term (based on adjusted p values (log q)).

## Results

### Divergence and plasticity of gene expression

Under common temperatures, heritable divergence in gene expression occurred between geothermal and ambient ecotypes (Figure 2A-B). Within the brain there were a greater number of divergently expressed genes between geothermal and ambient ecotypes at 12°C than at 18 °C (Figure 2; Table S1). Two genes were divergent between geothermal and ambient ecotypes regardless of temperature, *loc120815948* and *trpc4a*. In the liver, no genes were divergently expressed at either temperature (Figure S1; Table S1).

We detected plastic responses across both 12°C and 18°C in ecotypes and hybrids in the brain (Figure 3A-C; Table S2). The geothermal ecotype and hybrids showed a greater number of plastic genes compared to the ambient ecotype (Table S2). There were 47 shared plastic genes overlapping between the geothermal and ambient ecotypes (Table S2), accounting for 77% of the plasticity genes exhibited by the ambient ecotype. A similar proportion (70%) of plastic genes overlapped between the ambient ecotype and the hybrids. While the geothermal ecotype and hybrids shared 85 genes, the majority of their plastic responses were in ecotype-specific genes. Across ecotypes and hybrids there were 38 shared plastic responses showing evidence for enrichment of RNA processing (6 genes) and splicing (5 genes). Comparatively, there was a lower degree of plasticity in gene expression across 12°C and 18°C for ecotypes and hybrids in the liver relative to the brain (Figure 2SA-C; Table S2), with only one down-regulated gene in the liver, *ptpn6*.

**Figure 3.**
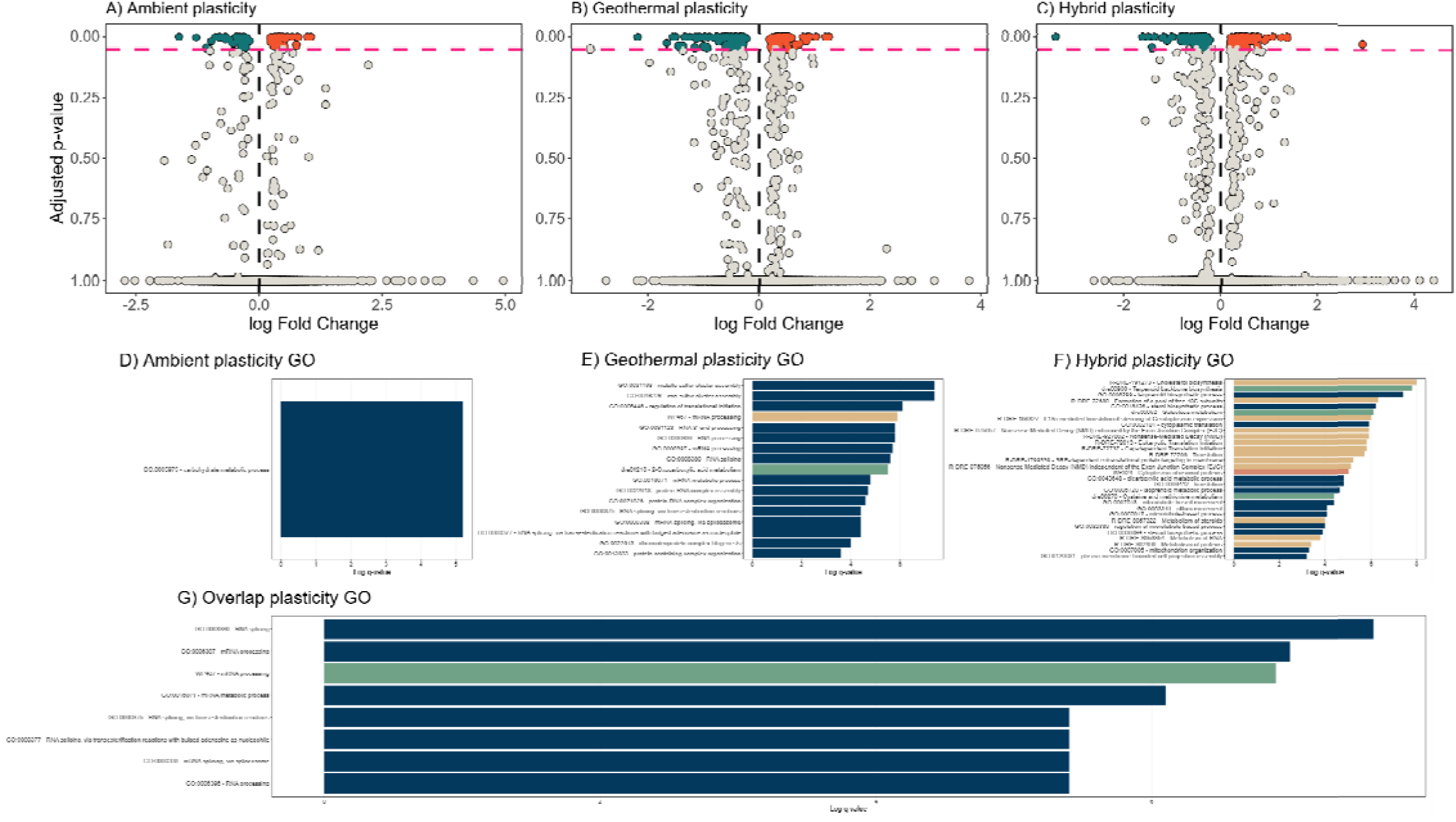
Volcano plots representing up- and down-regulated genes corresponding to plastic responses between rearing temperatures for ambient (A), geothermal (B), and hybrid (C) ecotypes of Icelandic stickleback. A-C) all up- and down-regulated genes are shown as orange and blue, respectively. The dashed red line highlights the p-value cut-off of 0.05. D-F) Gene ontology results highlighting the gene ontology terms from unique plastic responses to 12°C and 18°C rearing conditions for the ambient (D), geothermal (F), and hybrid (G) ecotype. G) gene ontology terms associated with the 38 genes showing common plastic responses across all three ecotypes. The different colours in each GO plot represent the database the term belongs to.

### Effects of hybridization on divergence and plasticity of gene expression

Divergence in gene expression for both ambient and geothermal ecotypes relative to hybrids was widespread in the brain (Figure 4A-D; Table S3). In total, 2045 genes were differentially expressed between the ambient ecotype and hybrids across rearing temperatures, with 553 and 531 genes being uniquely divergent at 12°C and 18°C repectively. A large proportion (78% at 12°C and 79% at 18°C) of differentially expressed genes were divergent at both temperatures between the ambient ecotype and hybrids indicating a strong heritable component. A similar pattern of divergence was observed between the geothermal ecotype and hybrids, with 1559 divergent genes across both rearing temperatures (78% at 12°C and 75% at 18°C), again indicating a strong heritable component of variation. There was a high degree of divergence in gene expression between both the ambient and geothermal ecotypes at 12°C and 18°C (Table S3). A total of 1395 genes were divergent between hybrids and both ecotypes regardless of rearing temperature. In contrast, only a small number of genes involved in expression divergence between hybrids and both ecotypes within the liver (Figure S4A-D; Table S3). There were only 2 divergent genes at 12°C and no divergent genes at 18°C between ecotypes relative to hybrids, with no overlap between comparisons observed.

**Figure 4.**
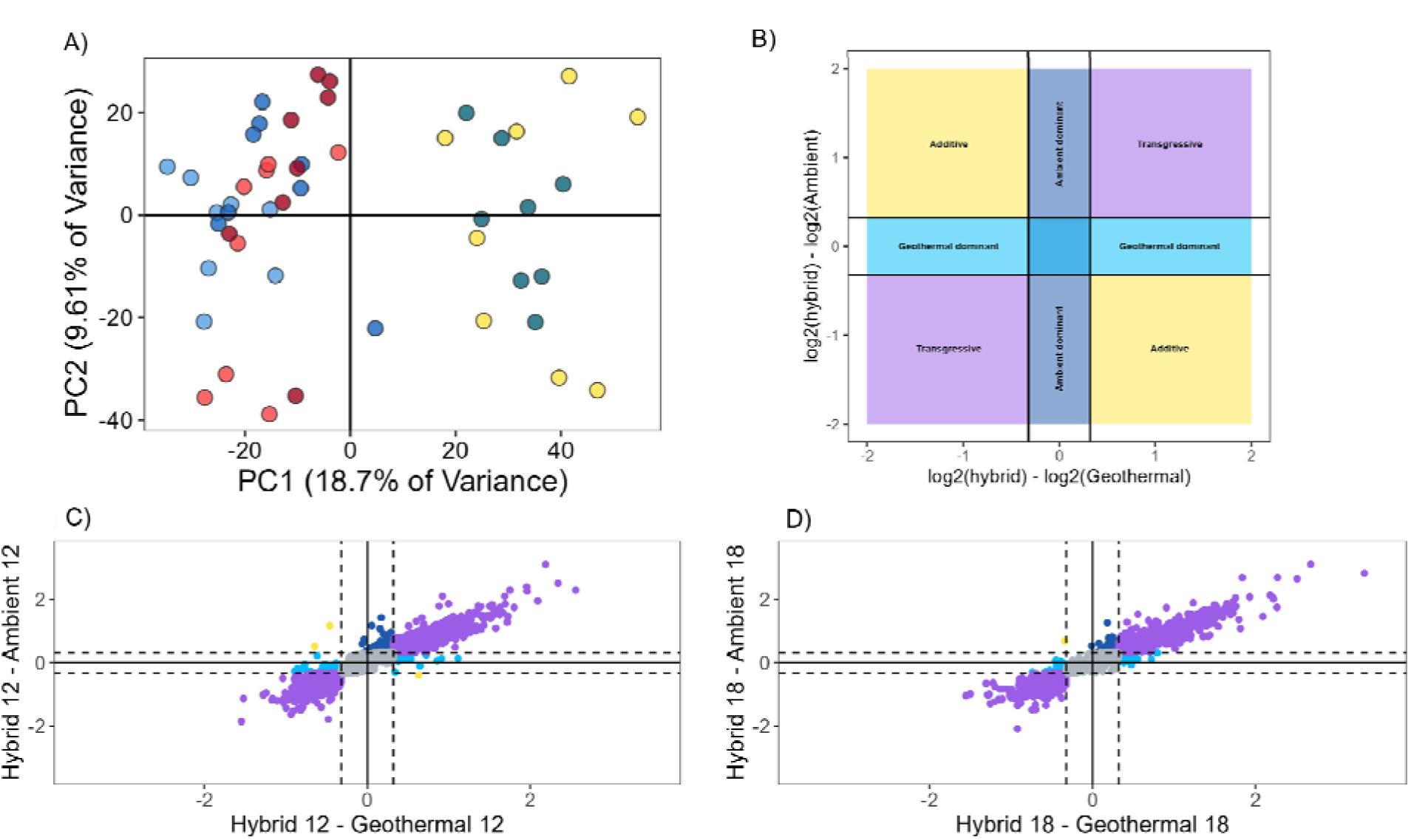
Inheritance patterns of divergent genes in hybrids compared to pure-strain geothermal and ambient ecotypes. A) Principal component analysis of all differentially expressed genes. Dark and light blue corresponds to the ambient ecotype at 12°C and 18°C. Dark and light red corresponds to the geothermal ecotype at 12°C and 18°C. Green and yellow corresponds to the hybrid ecotype at 12°C and 18°C. B) Patterns of inheritance based on log fold change between hybrids relative to each pure-strained ecotype. Dominant inheritance (light and dark blue) patterns are between +/- 0.32 for each comparison. Additive inheritance (yellow) is defined as >0.32 in geothermal but <0.32 in the ambient ecotype or <0.32 in the geothermal ecotype but >0.32 in the ambient ecotype. Transgressive inheritance (purple) is defined as either < or > 0.32 in each of the pure-strained ecotypes. C) Patterns of inheritance in all differentially expressed genes at 12°C. D) Patterns of inheritance in all differentially expressed genes at 18°C.

**Figure 5.**
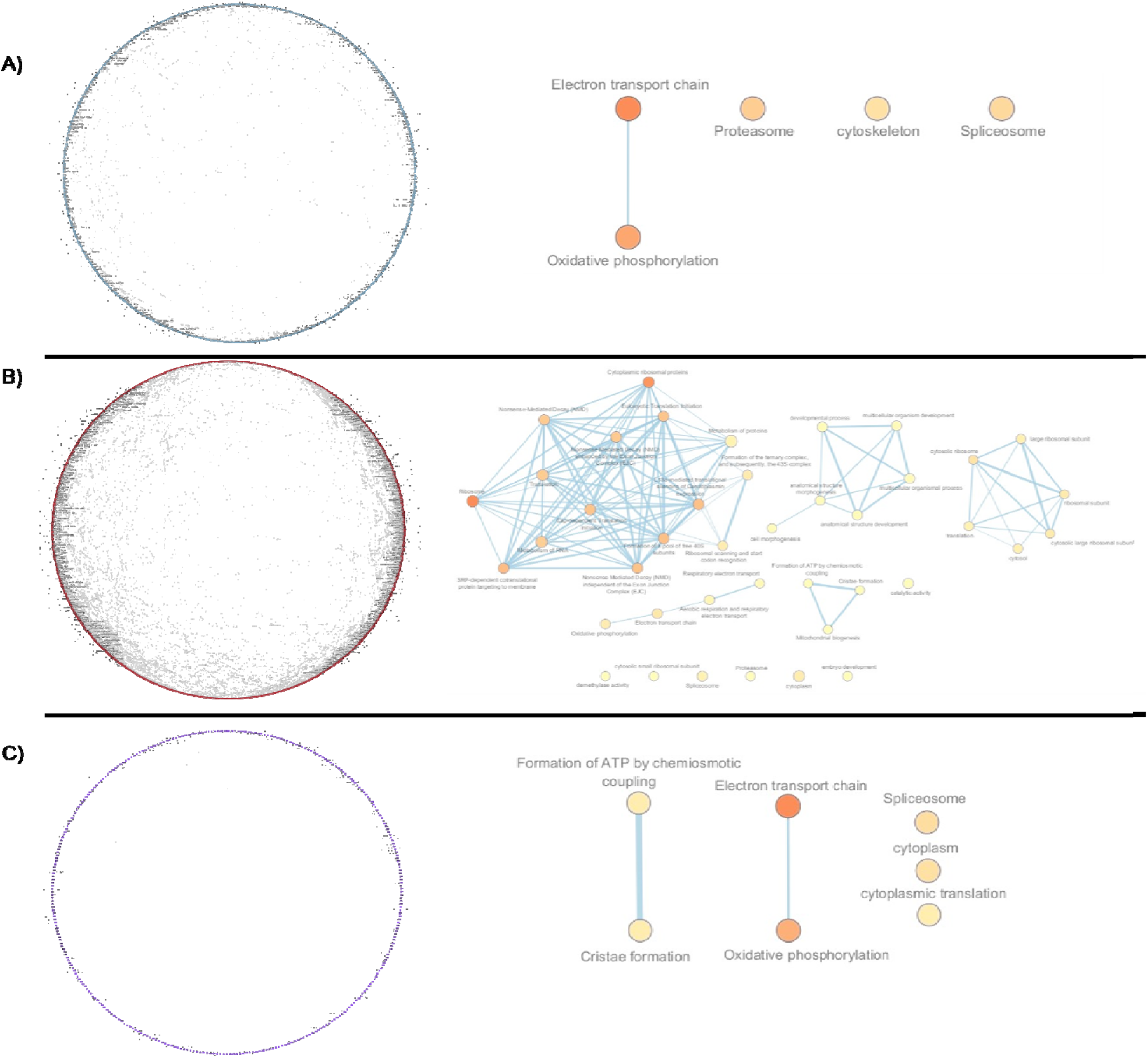
Gene network analyses for transgressively inherited genes in hybrids relative to pure-strained ecotypes. A) Gene network for transgressively inherited genes at 12°C and the significant gene ontology terms for the network. B) Gene network for transgressively inherited genes at 18°C and the corresponding gene ontology terms for the network. C) Overlap between the 12°C and 18°C network of transgressively inherited genes and the associated gene ontology terms for the overlapping network.

Regardless of temperature, there was substantial divergence of transgressively expressed genes between the hybrids and both ecotypes in the brain (Figure 4; Table S4). Hybrids were also clearly divergent from both ecotypes within the PCA (Figure 4A) with 87% of differentially expressed genes at 12°C and 18°C classified as transgressively expressed (Figure 4B-D; Table S4). Divergence between hybrids and ecotypes displayed 2405 transgressively expressed genes irrespective of rearing temperature. Of these, 354 were unique to 12°C, while 377 were unique to 18°C. In the liver, there were 15 transgressively expressed genes at 12°C and 4 at 18°C, with only 2 genes overlapping between temperatures.

### Gene co-expression networks

In the brain, whole gene networks were associated with plastic responses for each of the ambient and geothermal ecotypes and their hybrids (Figure S5A-C). There were 49 genes connected by 140 edges making up the gene network of plastic responses for the ambient ecotype (Figure S5A) with gene ontologies (GO) associated with DNA binding and mitochondrial ABC transporters (Table S5). Geothermal plastic response gene networks had 181 genes connected by 507 edges (Figure S5B) with GO terms associated with mRNA processing, spliceosome, defense responses, hydrolase activity, and cytoplasm (Table S5). Lastly, the gene network of hybrid plastic responses had 160 genes connected by 461 edges (Figure S5C) associated with the cytoplasm (Table S5). These three gene networks overlapped at 23 genes, connected by 8 edges, corresponding to lysozyme activity, CD4 receptor binding, and peptidoglycan muralytic activity (Table S5). All overlapping plastic gene networks appear to function as part of the immune system responses to increased temperature.

Divergence between ecotypes was dependent on temperature with expression at 12°C associated with a whole network of genes, but this was not the case at 18°C. The gene network associated with divergence at 12°C had 13 genes and 20 edges connecting the genes (Figure S6). Gene ontology analysis of the network identified *yrdc* as being associated with RNA binding and translational fidelity and *mgst3a* being involved in glutathione-dependent detoxification of reactive toxins. At 18°C, no genes had an association above 0.7 and a network could not be built.

Gene networks associated with divergence between ecotypes and hybrids contained more genes than any other network (Figure S7A-D), regardless of rearing temperature. These gene networks were extensive and contained 2470 genes with 4092 edges at 12°C and 2468 genes with 4147 edges at 18°C (Figure S7A-B) with a considerable number of overlapping genes (1945) and edges (1332) across rearing temperatures (78% overlap in genes at 12°C and 18°C). The gene networks associated with divergence between the geothermal ecotype and hybrids were slightly less extensive containing 1895 genes and 3113 edges at 12°C and 1986 genes and 3327 edges at 18°C. Overlap was similar as to ambient ecotypes with 1480 genes and 973 edges that were concordant between rearing temperatures (78% overlap at 12°C and 74% overlap at 18°C) and associated with divergence between geothermal ecotypes and hybrids. Across all divergence networks there were 1292 genes and 497 edges in common, but a relatively higher proportion of overlapping gene networks associated with geothermal-hybrid divergence (68.2% at 12°C and 65.1% at 18°C) than ambient-hybrid divergence (52.3% at 12°C and 50.3% at 18°C). This overlapping network was overall less connected, measured by overlapping edges, than the networks associated with ambient-hybrid divergence (12.1% overlap at 12°C and 12.0% overlap at 18°C) and geothermal-hybrid divergence (16.% overlap at 12°C and 14.9% overlap at 18°C).

We further identified gene networks associated with transgressive inheritance patterns and their biological function, specifically involving genes that were over- or under-expressed in hybrids relative to ecotypes. The gene network associated with transgressive expression in hybrids relative to the ambient ecotype had 1142 genes with 3242 edges at 12°C but was plastic by increasing to 1683 genes with 4762 edges when raised under 18°C. The geothermal ecotype showed a less extensive network associated with transgressive expression in hybrids with 765 genes with 2157 edges at 12°C but this increased substantially to 1320 genes with 3757 edges at 18°C. For both ecotypes there was overlap with 12°C conditions providing a gene network associated with transgressive expression having 612 genes with 674 edges and GO terms relating to the electron transport chain, oxidative phosphorylation, the cytoskeleton, the proteasome, and the spliceosome (Figure 6A; Table S7). The overlapping transgressive gene network at 18°C contained 1180 genes with 1481 edges (Figure 6B) with 42 significant GO terms primarily corresponding to mitochondrial function, energy production, and use (Table S7). Across both temperatures, the transgressive expression network contained 331 genes, 136 edges, and 7 GO terms related to energy production (Figure 6C; Table S7).

### Proportion of inheritance pattern of transgressive loci

Allele-specific expression (ASE) was detected in a small number of SNPs, indicating cis-regulatory mechanisms for four specific genes that overlapped across comparisons. Overall, SNPs within mfn1b, emsy, eef1a1, and fahd1 were identified as being under cis-regulatory control and associated with temperature-dependent expression differences between ecotypes. Three SNPs located in mfn1b, emsy, and eef1a1 were inherited from the ambient ecotype, whereas seven SNPs distributed across all four genes were inherited from the geothermal ecotype. ASE identified differential regulation among these loci where SNPs associated with eef1a1 and fahd1 exhibited reduced expression, while alleles at the remaining loci showed increased expression. The magnitude of these expression changes was temperature dependent, with larger differences observed at 12°C compared to 18°C, indicating that cid-regulatory variants at multiple loci contribute to temperature-associated divergence.

## Discussion

We investigated the genomic consequences of hybridization between locally adapted thermal ecotypes of Icelandic stickleback using a two temperature common garden experiment. First, we tested for evidence of expression divergence (including plasticity) by comparing geothermal and ambient ecotypes across temperatures. While we identified a low degree of gene expression divergence between ecotypes, many genes were plastic to thermal variation in ways unique to each ecotype. Second, we tested whether hybrid gene expression differed from ecotypes. Supporting this, hybrid gene expression was highly divergent from both ecotypes regardless of developmental temperature indicating strong heritable effects. The majority of divergent genes exhibited transgressive expression and suggested substantial disruptions occurring within gene networks, with increasing effects under the warmer condition. This suggested that hybridization between ecotypes incurs asymmetric effects across temperatures, with gene networks of transgressive expression primarily associated with energy use and recruitment (Barreto et al., 2015a; Moran et al., 2021; Schwartz et al., 2024). Further, warmer environmental conditions disrupted a greater number of functional pathways suggesting maladaptive phenotypes may be more likely to arise. Third, we tested if hybridization between locally adapted ecotypes disrupted co-adapted cis-trans regulatory networks, leading to misexpression of specific alleles of locally adapted genes. Counter to our hypothesis, we found a surprisingly low degree of transgressively expressed SNPs within genes which suggests a greater degree of trans- vs cis-regulatory changes in gene expression due to hybridization. However, genes associated with cis-regulatory changes were associated with appear to have effects on DNA repair (*emsy*), metabolism and heat shock response (*eef1a1*), and mitochondrial function (*mfn1b* and *fahd1*). Thus, even though a low degree of genomic divergence occurred between ecotypes (Brachmann et al., 2025), the impacts of hybridization could involve metabolic dysfunction that may incur severe environmentally dependent fitness consequences.

### Divergence and plasticity of thermal ecotypes

The low degree of gene expression divergence between ecotypes could be due to the rapid pace of evolution found in this system. The population pair used here was established only ∼70 years ago by the output of geothermal heating systems in nearby residential buildings. Despite the geothermal environment being colonized recently, and low levels of genome-wide allelic differentiation from ambient counterparts (Brachmann et al., 2025), the gene expression patterns disrupted by hybridization were key for energy use and metabolism. Such principles could underlie other examples of rapid evolutionary divergence (Hairston et al., 2005; Kopp & Matuszewski, 2014; Messer et al., 2016), including other cases of fish adapting to warm habitats (Dayan et al., 2019). Our findings suggest that despite low levels of expression divergence reproductive isolation could be enhanced by plasticity, facilitating and maintaining phenotypic divergence between thermal habitats.

While there were low levels of divergence in gene expression, plasticity was divergent. Ecotypes, as well as hybrids, showed unique plastic responses to temperature, with geothermal ecotypes and hybrids showing a greater number of plastic genes relative to the ambient ecotype. Overall, the plastic responses of the ambient ecotype to warmer conditions were associated with carbohydrate metabolism, indicating plastic shifts in the energy supply required in warming environments (Madeira et al., 2017; Middleton et al., 2024; Moore et al., 2025). In contrast, geothermal ecotypes appear to adapt to warming environments through the tuning of alternative splicing regulatory mechanisms (Schlaen et al., 2015; Steward et al., 2022; Haltenhof et al., 2024) and overall metabolic architecture to manage chronic metabolic and oxidative challenges (Seebacher et al., 2010; Madeira et al., 2016; Lemieux & Blier, 2022). Divergent plastic responses, indicative of adaptation to warming, is thus facilitated through different metabolic strategies. In contrast, hybrids had relatively biologically broad plastic responses suggestive of a release of unrefined cryptic genetic variation (Gibson & Dworkin, 2004; Vallejo-Marín & Hiscock, 2016; Mikkelsen & Irwin, 2021) that may be maladaptive in both environments. The divergence in the plasticity of metabolic processes corresponds with our previous findings of ecotype differences in metabolic rate (Pilakouta et al., 2020), suggesting that hybridization may drive maladaptive plasticity in metabolic rate (Ghalamboret al., 2007; Chevin & Hoffmann, 2017; Campbell-Staton et al., 2021; Eriksson et al., 2023).

While ecotype specific plastic responses to thermal variation were identified, we also detected shared plastic responses across ecotypes. For example, hepatic *ptptn6* showed a shared plastic response to thermal variation. *Ptpn6* is a known regulator of the immune system through the *shp1* protein (Kanwal et al., 2013) and has been implicated in regulating insulin resistance and lipid metabolism (Marín-Juez et al., 2014; Xu et al., 2014, 2014; Kumar et al., 2023). In zebrafish (*Dania rerio*), *ptpn6/shp1* have previously been shown to play a critical role in modulating insulin-resistant and insulin sensitive states (Marín-Juez et al., 2014). The ambient ecotype showed greater downregulation of ptpn6 in response to warmer temperatures than the geothermal ecotype or hybrids, suggesting a potentially stronger compensatory response that may mitigate the downregulation of insulin signaling and immune pathways under warmer developmental environments. In the brain, genes showing common plastic responses (the same gene and direction of plastic response) comprised 62% of the total amount of plastic genes in the ambient ecotype compared to only 16% and 20% in the geothermal ecotype and hybrids. The high proportion of common plastic responses in the ambient ecotype indicated conserved and possible ancestral plastic responses (Wund et al., 2008; Wood et al., 2023; Coates et al., 2025), while the low proportion of common plastic responses in the geothermal ecotype suggested a more derived and ecotype-specific responses due to warming environments. The hybrids did not appear to combine parental plastic responses which may indicate potential regulatory incompatibilities between ecotypes (Go & Civetta, 2020; Lemmon & Kirkpatrick, 2006; Vallejo-Marín & Hiscock, 2016).

### Effects of hybridization on gene expression

Selection against hybridization can contribute to the formation and maintenance of reproductive isolation between diverging populations (Butlin et al., 2014; Kulmuni et al., 2020). Divergent populations are expected to be locally adapted and hybrids inheriting intermediate or transgressive phenotypes are likely to face fitness consequences (Chhina et al., 2022; Gow et al., 2007; Hartman et al., 2013; Rice & McQuillan, 2018; Thompson et al., 2021). As ecotype specific gene expression is expected to be adapted to their native habitats (Ayroles et al., 2009; Brauer et al., 2017), uncoordinated gene expression or misexpression in hybrids is likely to be maladaptive in the wild. Hybrid gene expression differed greatly from both geothermal and ambient ecotypes at both rearing temperatures and predominantly showed transgressive patterns of inheritance. Hybrid mis-expression has been found to accompany severe (Barreto et al., 2015b; Ellison & Burton, 2008; Ortíz-Barrientos et al., 2007) or unpredictable (Brice et al., 2021; Díaz et al., 2023) fitness consequences (Rogers & Bernatchez, 2006; Rogers et al., 2007). Here, the disruption of gene co-expression networks was enhanced by the warmer temperature, suggesting thermally-dependent regulatory incompatibilities between ecotypes. Similar findings for hybrids has been shown in Atlantic silversides (*Menidia menidia*) (Jacobs et al., 2024), where a high proportion of genes were mis-expressed in relative to parental ecotypes. In Icelandic stickleback, transgressive mis-expression appeared to disrupt metabolic processes for energy use and production, consistent with hybrid metabolic dysfunction (Barreto et al., 2015). Our data indicate that the breakdown of metabolic processes in hybrids could be underlain by mitochondrial dysfunction, representing a severe physiological consequence (Barreto et al., 2015). Finally, the consequences of hybridization were enhanced in the warmer condition, further supporting the role of thermal plasticity in promoting reproductive isolation, although asymmetrically between thermal habitats.

### Regulatory mechanisms and modes of inheritance underlying divergence

We found little evidence that additive variation contributed to divergence between geothermal and ambient sticklebacks. While transcriptional variation has been shown to be largely additive (Gilad et al., 2008; Kim & Gibson, 2010), our findings indicated less than 1% of such variation was additive across temperatures. While this would classically be interpreted as suggesting little potential for evolution (Falconer & Mackay, 1996), widespread non-additive gene expression inheritance has previously been found in other vertebrates (Debes et al., 2012), including stickleback (Leder et al., 2014). Hybridization transmitted a high degree of non-additive transcriptional variation indicating complex interactions due to dominance, epistasis, and transgressive variation (Yazdi et al., 2022; Jacobs et al., 2024; Tsouris et al., 2024). High levels of non-additive transcriptional variation is known to occur in complex phenotypic traits, such as metabolic phenotypes (Zhong et al., 2019; Zhang et al., 2025). However, a limitation of this study is that hybrid crosses were generated from a single population of divergent ecotypes with no reciprocal crosses. Future work using a fully reciprocal cross with multiple divergent populations would allow for a more revealing examination of additive and non-additive transcriptional variation.

Transgressive expression of genes in hybrids, relative to either parental ecotype, was the dominant pattern of gene expression at both rearing temperatures, suggesting substantial disruption of regulatory mechanisms. Generally, cis-regulatory mechanisms are expected to underlie evolutionary divergence in gene expression more often than trans-regulatory changes (Signor & Nuzhdin, 2018; Verta & Jones, 2019; Wittkopp et al., 2004; Zhong et al., 2019). However, we identified a low degree of cis-regulation and a high degree of trans-regulation of transgressively expressed genes associated with metabolic divergence. Trans-regulation has previously been shown to govern non-additive transcriptional variation (Tsouris et al., 2024) and may play a large role temperature dependent regulation of metabolic phenotypes as they rapidly respond to environmental variation (Chen & Zhu, 2004).

Hybridization between locally adapted and rapidly evolving thermal ecotypes appears to drive genomic incompatibilities and maladaptive phenotypic variation, particularly under elevated temperatures. These findings have important implications for climate change biology. As global temperatures rise, thermal gradients are expected to shift, potentially increasing contact between locally adapted populations that were previously segregated by microhabitat differences. Hybridization is often considered a mechanism for evolutionary rescue and increased adaptive potential under environmental change (Kelly & Phillips, 2016; Pfennig et al., 2016; Kulmuni et al., 2023; Sexton et al., 2024). In contrast, our results demonstrated that even small amounts of genomic divergence can generate temperature-dependent incompatibilities which may cause reduced fitness. In this system, warmer environments intensified regulatory disruption, suggesting that climate warming may not only impose direct physiological stress but also destabilize gene regulatory networks shaped by local adaptation. These findings illustrate that population mixing or assisted gene flow may dampen resilience to warming environments. Instead, the outcome of hybridization may critically depend on the degree of regulatory co-adaptation and the environmental context in which hybrids develop. As anthropogenic warming continues to alter thermal landscapes, understanding when hybridization facilitates adaptation versus when it uncovers cryptic incompatibilities will be essential for predicting evolutionary responses and guiding conservation strategies.

## Supporting information

All supplementary figures

all supplementary tables

