## Supplementary material for "Energetic misfires: Hybridization drives transgressive expression in metabolic pathways in thermally divergent Icelandic stickleback": All supplementary figures

**Table S1.** Differentially expressed genes in brain and liver tissue between geothermal and ambient ecotypes raised under 12°C and 18°C.

| Tissue | Temperature | DE genes | Up-regulated | Down-regulated |
|---|---|---|---|---|
| Brain | 12°C | 18 | 11 | 7 |
| Brain | 18°C | 8 | 3 | 5 |
| Liver | 12°C | 0 | 0 | 0 |
| Liver | 18°C | 0 | 0 | 0 |

**Table S2.** Differentially expressed genes between 12 and 18°C for geothermal and ambient ecotypes and their hybrids for brain and liver tissue. Each differentially expressed gene changes in log-fold change between rearing environments, indicating plasticity.

| Ecotype | Tissue | DE genes | Up-regulated | Down-regulated |
|---|---|---|---|---|
| Geothermal | Brain | 202 | 116 | 86 |
| Ambient | Brain | 61 | 32 | 29 |
| Hybrid | Brain | 189 | 87 | 102 |
| Geothermal-Ambient overlap | Brain | 47 | 26 | 21 |
| Geothermal-hybrid overlap | Brain | 85 | 46 | 39 |
| Ambient-hybrid overlap | Brain | 43 | 22 | 21 |
| Complete overlap | Brain | 38 | 20 | 18 |
| Geothermal | Liver | 3 | 1 | 2 |
| Ambient | Liver | 4 | 2 | 2 |
| Hybrid | Liver | 5 | 3 | 2 |
| Geothermal-Ambient overlap | Liver | 1 | 0 | 1 |
| Geothermal-hybrid overlap | Liver | 1 | 0 | 1 |
| Ambient-hybrid overlap | Liver | 2 | 1 | 1 |
| Complete overlap | Liver | 1 | 0 | 1 |

**Table S3.** Differentially expressed genes associated with divergence between thermal ecotypes and hybrids in 12 and 18°C rearing temperatures for both brain and liver tissue. Overlap indicates differentially expressed genes overlapping between 12 and 18°C. Pure indicates overlap between geothermal and ambient ecotypes.

| Comparison | Tissue | Type | Temperature | DE genes | Up-regulated | Down-regulated |
|---|---|---|---|---|---|---|
| Geothermal-hybrid | Brain | Divergence | 12°C | 1994 | 1218 | 776 |
| Geothermal-hybrid | Brain | Divergence | 18°C | 2081 | 1159 | 922 |
| Geothermal-hybrid | Brain | Divergence | Overlap | 1559 | 933 | 626 |
| Ambient-hybrid | Brain | Divergence | 12°C | 2598 | 1513 | 1085 |
| Ambient-hybrid | Brain | Divergence | 18°C | 2576 | 1469 | 1107 |
| Ambient-hybrid | Brain | Divergence | Overlap | 2045 | 1208 | 837 |
| Pure-hybrid | Brain | Divergence | 12°C | 1763 | 1094 | 669 |
| Pure-hybrid | Brain | Divergence | 18°C | 1815 | 1041 | 774 |
| Pure-hybrid | Brain | Divergence | Overlap | 1395 | 852 | 543 |
| Geothermal-hybrid | Liver | Divergence | 12°C | 2 | 1 | 1 |
| Geothermal-hybrid | Liver | Divergence | 18°C | 0 | 0 | 0 |
| Geothermal-hybrid | Liver | Divergence | Overlap | 0 | 0 | 0 |
| Ambient-hybrid | Liver | Divergence | 12°C | 2 | 0 | 2 |
| Ambient-hybrid | Liver | Divergence | 18°C | 0 | 0 | 0 |
| Ambient-hybrid | Liver | Divergence | Overlap | 0 | 0 | 0 |
| Pure-hybrid | Liver | Divergence | 12°C | 0 | 0 | 0 |
| Pure-hybrid | Liver | Divergence | 18°C | 0 | 0 | 0 |
| Pure-hybrid | Liver | Divergence | Overlap | 0 | 0 | 0 |

| Brain | | | Liver | | |
|---|---|---|---|---|---|
| Type | Temperature | DE genes | Type | Temperature | DE genes |
| Dominant geothermal | 12°C | 170 | Dominant geothermal | 12°C | 4 |
| Dominant ambient | 12°C | 237 | Dominant ambient | 12°C | 3 |
| Additive | 12°C | 3 | Additive | 12°C | 1 |
| Transgressive | 12°C | 2759 | Transgressive | 12°C | 15 |
| Dominant geothermal | 18°C | 184 | Dominant geothermal | 18°C | 4 |
| Dominant ambient | 18°C | 238 | Dominant ambient | 18°C | 8 |
| Additive | 18°C | 1 | Additive | 18°C | 2 |
| Transgressive | 18°C | 2782 | Transgressive | 18°C | 4 |

**Table S4.**

Inheritance patterns of differentially expressed genes at 12 and 18°C for both brain and liver tissues.

**Table S5.** Gene ontology of transgressive gene network expression associated with plastic responses between 12°C and 18°C for thermal ecotypes and hybrids.

| Comparison | GO description | GO Terms | Genes | P-value |
|---|---|---|---|---|
| Geothermal plasticity | mRNA processing | WP:WP467 | fkbp3\|rtca\|pes\|ENSGACG00000007556\|si:dkey-19e4.5\|ENSGACG00000012425\|med13b\|cct4 | 1.4120471366123126E-5 |
| Geothermal plasticity | Spliceosome | KEGG:03040 | rtca\|pes\|ENSGACG00000007556\|rab11al\|si:dkey-19e4.5\|ENSGACG00000012425\|med13b\|cct4\|sirt7 | 3.406732128265136E-5 |
| Geothermal plasticity | defense response to bacterium | GO:0042742 | snrpa\|SLC1A3\|col4a3 | 0.004727 |
| Geothermal plasticity | hydrolase activity | GO:0016811 | paqr3b\|CDK19 | 0.020102392317369263 |
| Geothermal plasticity | cytoplasm | GO:0005737 | mob4\|dhx15\|lrrc45\|ENSGACG00000013294\|paqr3b\|hcfc1a\|wasf3b\|ENSGACG00000000821\|NFE2L1\|col4a3 | 0.046715755632638176 |
| Ambient plasticity | DNA binding | GO:0008301 | *ensgacg00000020324\| ints8* | 0.003338 |
| Ambient plasticity | Mitochondrial ABC transporters | REAC:R-DRE-1369007 | Isyna1 | 0.049939 |
| Hybrid plasticity | cytoplasm | GO:0005737 | ciarta\|ypel3\|enkur\|stk17b\|tbcelb\|acy1\|rpl23a\|ENSGACG00000014965\|stt3b\|tppp3 | 0.002468 |

| | | | | |
|---|---|---|---|---|
| Overlap | lysozyme activity | GO:0003796 | ENSGACG00000014965 | 0.014948256036796614 |
| Overlap | CD4 receptor binding | GO:0042609 | mapre3b | 0.014948256036796614 |
| Overlap | peptidoglycan muralytic activity | GO:0061783 | ENSGACG00000014965 | 0.029890781478982165 |

**Table S6.** Divergence between geothermal and ambient ecotype per temperature - gene networks

| Temperature | GO description | GO Terms | Genes | P-value |
|---|---|---|---|---|
| 12 | regulation of translational fidelity | GO:0006450 | yrdc | 0.001725 |
| 12 | Aflatoxin activation and detoxification | REAC:R-DRE-5423646 | mgst3a | 0.012561060711794081 |
| 12 | tRNA binding | GO:0000049 | yrdc | 0.016864699118435578 |
| 12 | double-stranded RNA binding | GO:0003725 | yrdc | 0.016864699118435578 |
| 12 | Glutathione conjugation | REAC:R-DRE-156590 | mgst3a | 0.035589672016748786 |
| 12 | nucleotidyltransferase activity | GO:0016779 | yrdc | 0.046378 |

**Table S7.** Gene ontology of transgressively expressed gene networks diverging between both geothermal and ambient ecotypes and hybrids.

| Temperature | GO description | GO Terms | Genes | P-value |
|---|---|---|---|---|
| 12 | Oxidative phosphorylation | KEGG:00190 | pop4\|eif4g3a\|ache\|erbin\|ankib1b\|tcta\|ints6l\|gnsb\|ENSGACG00000010347\|sprb\|zgc:85722\|rhebl1\|tanc2a\|ttc21b\|tmem147\|MDGA2\|ccdc186\|trioa | 3.7228208859649733E-7 |
| 12 | Electron transport chain | WP:WP1339 | ogal\|ache\|ankib1b\|tcta\|ints6l\|gnsb\|ENSGACG00000010347\|sprb\|zgc:85722\|rhebl1\|tanc2a\|MDGA2\|ccdc186\|trioa | 3.0311642466659246E-6 |
| 12 | Oxidative phosphorylation | WP:WP1335 | eif4g3a\|ache\|WRB\|ankib1b\|tcta\|ENSGACG00000010347\|rhebl1\|tanc2a\|MDGA2\|trioa | 5.3127893323819846E-5 |
| 12 | Proteasome | KEGG:03050 | rbm45\|lsm4\|bccip\|timm17a\|ENSGACG00000020674\|ENSGACG00000006143\|naa25\|nbr1b | 0.00389 |
| 12 | Spliceosome | KEGG:03040 | pnpo\|rhbdf1a\|ENSGACG00000007877\|ctcf\|cdc42l\|rnls\|pdss2\|ikbkg\|ube2a\|ENSGACG00000019219\|ENSGACG00000001206\|tcf3b | 0.015004040528928486 |
| 12 | cytoskeleton | GO:0005856 | ENSGACG00000019647\|heatr6\|csnk1g1\|SNX14\|ENSGACG00000011427\|citb\|sclt1\|SNRPD1\|afap1 | 0.0407 |
| 18 | Ribosome | KEGG:03010 | top2b\|chia.1\|plxna4\|rangrf\|surf1\|ENSGACG00000007324\|ablim3\|gstr\|sgsm1a\|si:d | 7.208875004358157E-52 |

| | | | | |
|---|---|---|---|---|
| | | | key-119f1.1\|FUCA1\|FAM135B\|lrba\|rps28\|ENSGACG00000017543\|smpdl3a\|ctbs\|KCNMA1\|mrpl41\|zbtb8a\|znf609b\|lancl1\|etfb\|rnasekb\|ciapin1\|gnb5b\|ENSGACG00000007357\|si:ch211-14a17.11\|NDUFB1\|smim15\|chmp3\|gstk4\|usp38\|usp32\|march7\|NDUFA4\|gm2a\|ndufb10\|micu3a\|usp20\|fynb\|wdr60\|fam32a\|ano11\|ptgesl\|psma3\|tiprl\|lyrm7\|ANO4\|tecpr2\|rpl39\|rpl30\|rpl31\|mrps10\|psmd6\|ENSGACG00000009156\|arl6ip4\|cnot1\|ENSGACG00000016782\|si:dkey-266j7.2\|grk4\|ENSGACG00000014564\|galm\|ENSGACG00000015058\|ENSGACG00000011934 | |
| 18 | Cytoplasmic ribosomal proteins | WP:WP324 | rangrf\|surf1\|ENSGACG00000007324\|ablim3\|gstr\|sgsm1a\|si:dkey-119f1.1\|FUCA1\|FAM135B\|rps28\|aclyb\|KCNMA1\|zbtb8a\|znf609b\|lancl1\|etfb\|rnasekb\|ciapin1\|gnb5b\|ENSGACG00000007357\|si:ch211-14a17.11\|NDUFB1\|smim15\|chmp3\|gstk4\|usp38\|usp32\|march7\|NDUFA4\|gm2a\|micu3a\|usp20\|fynb\|wdr60\|fam32a\|ano11\|psma3\|ANO4\|tecpr2\|rpl39\|rpl30\|rpl31\|mrps10\|psmd6\|ENSGACG00000009156\|arl6ip4\|cnot1\|ENSGACG00000016782\|si:dkey-266j7.2\|grk4\|ENSGACG00000014564\|galm\|ENSGACG00000015058\|ENSGACG00000011934 | 2.30371538629667E-47 |
| 18 | SRP-dependent cotranslational protein targeting to membrane | REAC:R-DRE-1799339 | nudt5\|rangrf\|surf1\|ENSGACG00000007324\|gstr\|sgsm1a\|si:dkey-119f1.1\|FUCA1\|FAM135B\|rps28\|KCNMA1\|etfb\|gnb5b\|ENSGACG00000007357\|si:ch211- | 2.6774243534507785E-30 |

| | | | | |
|---|---|---|---|---|
| | | | 14a17.11\|NDUFB1\|smim15\|chmp3\|gstk4\|usp32\|march7\|gm2a\|ndufb10\|micu3a\|usp20\|wdr60\|fam32a\|ano11\|psma3\|tecpr2\|rpl39\|rpl30\|rpl31\|mrps10\|psmd6\|ENSGACG00000009156\|arl6ip4\|cnot1\|si:dkey-266j7.2\|eif5b\|grk4\|ENSGACG00000014564\|ENSGACG00000011934 | |
| 18 | Formation of a pool of free 40S subunits | REAC:R-DRE-72689 | rangrf\|surf1\|ENSGACG00000007324\|gstr\|sgsm1a\|si:dkey-119f1.1\|FUCA1\|FAM135B\|rps28\|KCNMA1\|etfb\|gnb5b\|ENSGACG00000007357\|si:ch211-14a17.11\|NDUFB1\|smim15\|chmp3\|gstk4\|usp32\|march7\|gm2a\|ndufb10\|micu3a\|usp20\|wdr60\|fam32a\|ano11\|psma3\|psma5\|msra\|tecpr2\|rpl39\|rpl30\|rpl31\|mrps10\|psmd6\|ENSGACG00000009156\|arl6ip4\|cnot1\|si:dkey-266j7.2\|grk4\|ENSGACG00000014564\|ENSGACG00000011934 | 8.131549629163071E-29 |
| 18 | L13a-mediated translational silencing of Ceruloplasmin expression | REAC:R-DRE-156827 | eif2s3\|rangrf\|surf1\|ENSGACG00000007324\|gstr\|sgsm1a\|si:dkey-119f1.1\|FUCA1\|FAM135B\|rps28\|KCNMA1\|etfb\|gnb5b\|ENSGACG00000007357\|si:ch211-14a17.11\|NDUFB1\|smim15\|chmp3\|gstk4\|usp32\|march7\|gm2a\|ndufb10\|micu3a\|usp20\|wdr60\|fam32a\|ano11\|psma3\|psma5\|msra\|tecpr2\|rpl39\|rpl30\|rpl31\|mrps10\|psmd6\|ENSGACG00000009156\|arl6ip4\|cnot1\|si:dkey-266j7.2\|grk4\|ENSGACG00000014564\|ENSGACG00000011934 | 1.8145333361422437E-28 |
| 18 | Nonsense Mediated Decay (NMD) independent of | REAC:R-DRE-975956 | rangrf\|surf1\|ENSGACG00000007324\|gstr\|sgsm1a\|si:dkey-119f1.1\|FUCA1\|FAM135B\|r | 4.690302628290297E-27 |

| | | | | |
|---|---|---|---|---|
| | the Exon Junction Complex (EJC) | | ps28\|KCNMA1\|etfb\|gnb5b\|ENSGACG00000007357\|si:ch211-14a17.11\|NDUFB1\|smim15\|chmp3\|gstk4\|usp32\|march7\|gm2a\|ndufb10\|micu3a\|usp20\|wdr60\|fam32a\|ano11\|psma3\|tecpr2\|rpl39\|rpl30\|rpl31\|mrps10\|psmd6\|ENSGACG00000009156\|arl6ip4\|cnot1\|si:dkey-266j7.2\|grk4\|ENSGACG00000014564\|ENSGACG00000011934 | |
| 18 | Cap-dependent Translation Initiation | REAC:R-DRE-72737 | eif2s3\|rangrf\|surf1\|ENSGACG00000007324\|gstr\|sgsm1a\|si:dkey-119f1.1\|FUCA1\|FAM135B\|rps28\|KCNMA1\|etfb\|gnb5b\|ENSGACG00000007357\|si:ch211-14a17.11\|NDUFB1\|smim15\|chmp3\|gstk4\|usp32\|march7\|gm2a\|ndufb10\|micu3a\|usp20\|wdr60\|fam32a\|ano11\|psma3\|psma5\|msra\|tecpr2\|rpl39\|rpl30\|rpl31\|mrps10\|psmd6\|ENSGACG00000009156\|arl6ip4\|cnot1\|si:dkey-266j7.2\|grk4\|ENSGACG00000014564\|ENSGACG00000011934 | 5.870406681740524E-27 |
| 18 | Eukaryotic Translation Initiation | REAC:R-DRE-72613 | eif2s3\|rangrf\|surf1\|ENSGACG00000007324\|gstr\|sgsm1a\|si:dkey-119f1.1\|FUCA1\|FAM135B\|rps28\|KCNMA1\|etfb\|gnb5b\|ENSGACG00000007357\|si:ch211-14a17.11\|NDUFB1\|smim15\|chmp3\|gstk4\|usp32\|march7\|gm2a\|ndufb10\|micu3a\|usp20\|wdr60\|fam32a\|ano11\|psma3\|psma5\|msra\|tecpr2\|rpl39\|rpl30\|rpl31\|mrps10\|psmd6\|ENSGACG00000009156\|arl6ip4\|cnot1\|si:dkey-266j7.2\|grk4\|ENSGACG00000014564\|ENSGACG00000011934 | 1.0158016824388385E-26 |

| | | | | |
|---|---|---|---|---|
| 18 | Nonsense-Mediated Decay (NMD) | REAC:R-DRE-927802 | rangrf\|surf1\|ENSGACG00000007324\|gstr\|sgsm1a\|si:dkey-119f1.1\|FUCA1\|FAM135B\|rps28\|KCNMA1\|papolg\|etfb\|gnb5b\|ENSGACG00000007357\|si:ch211-14a17.11\|NDUFB1\|smim15\|chmp3\|gstk4\|usp32\|march7\|gm2a\|ndufb10\|micu3a\|usp20\|wdr60\|fam32a\|ano11\|psma3\|tecpr2\|rpl39\|rpl30\|rpl31\|mrps10\|psmd6\|ENSGACG00000009156\|arl6ip4\|cnot1\|si:dkey-266j7.2\|grk4\|ENSGACG00000014564\|ENSGACG00000011934 | 2.447850425333695E-24 |
| 18 | Nonsense Mediated Decay (NMD) enhanced by the Exon Junction Complex (EJC) | REAC:R-DRE-975957 | rangrf\|surf1\|ENSGACG00000007324\|gstr\|sgsm1a\|si:dkey-119f1.1\|FUCA1\|FAM135B\|rps28\|KCNMA1\|papolg\|etfb\|gnb5b\|ENSGACG00000007357\|si:ch211-14a17.11\|NDUFB1\|smim15\|chmp3\|gstk4\|usp32\|march7\|gm2a\|ndufb10\|micu3a\|usp20\|wdr60\|fam32a\|ano11\|psma3\|tecpr2\|rpl39\|rpl30\|rpl31\|mrps10\|psmd6\|ENSGACG00000009156\|arl6ip4\|cnot1\|si:dkey-266j7.2\|grk4\|ENSGACG00000014564\|ENSGACG00000011934 | 2.447850425333695E-24 |
| 18 | Translation | REAC:R-DRE-72766 | eif2s3\|top2b\|plxna4\|nudt5\|rangrf\|surf1\|ENSGACG00000007324\|gstr\|sgsm1a\|si:dkey-119f1.1\|FUCA1\|FAM135B\|rps28\|SUMF2\|KCNMA1\|etfb\|gnb5b\|ENSGACG00000007357\|si:ch211-14a17.11\|smim19\|NDUFB1\|smim15\|chmp3\|gstk4\|usp32\|march7\|emc10\|gm2a\|ndufb10\|micu3a\|usp20\|wdr60\|fam32a\|ano11\|psma3\|psma5\|msra\|tiprl\|tecpr2\|rpl39\|rpl30\|rpl31\|mrps10\|psmd6\|EN | 3.8103091718516755E-24 |

| | | | | |
|---|---|---|---|---|
| | | | SGACG00000009156\|arl6ip4\|cnot1\|si:dkey-266j7.2\|eif5b\|grk4\|ENSGACG00000014564\|ENSGACG00000011934 | |
| 18 | Metabolism of RNA | REAC:R-DRE-8953854 | nubp1\|ENSGACG00000015196\|rangrf\|surf1\|ENSGACG00000007324\|gstr\|sgsm1a\|si:dkey-119f1.1\|FUCA1\|FAM135B\|rps28\|KCNMA1\|tmem63c\|ENSGACG00000008852\|ccdc90b\|papolg\|etfb\|sec24c\|gnb5b\|ENSGACG00000007357\|si:ch211-14a17.11\|NDUFB1\|smim15\|chmp3\|gstk4\|usp32\|march7\|gm2a\|ndufb10\|micu3a\|usp20\|wdr60\|fam32a\|ano11\|psenen\|psma3\|mettl5\|tecpr2\|rpl39\|rpl30\|rpl31\|mrps10\|psmd6\|ENSGACG00000009156\|arl6ip4\|cnot1\|ENSGACG00000002373\|PPFIA3\|si:dkey-266j7.2\|timm8a\|grk4\|acad11\|ENSGACG00000014564\|ENSGACG00000011934\|magoh\|sqstm1\|aimp1\|ndufb6\|elp5 | 2.6322204084468037E-22 |
| 18 | Oxidative phosphorylation | KEGG:00190 | cbr1l\|sez6l2\|si:dkeyp-75h12.5\|imp3\|naa50\|lmbrd2b\|psmc3\|ENSGACG00000009179\|ENSGACG00000001473\|gpx4b\|eif4a1b\|tmco1\|rpl24\|rpl28\|rpl29\|mrps24\|rpl11\|ENSGACG00000017620\|TMEM250\|nbeaa\|fbl\|si:dkey-26c10.5\|fance\|lamtor4\|snrnp70\|atp5l\|mrpl3\|ENSGACG00000013492\|rpl7l1\|efhc1\|grnb\|ENSGACG00000010600\|clpp\|ogal\|rpl6\|tbc1d31\|ENSGACG00000003088\|gemin2 | 1.0522880680493652E-16 |
| 18 | cytosolic ribosome | GO:0022626 | gstr\|FUCA1\|zbtb8a\|usp32\|march7\|ndufb10\|fynb\|ANO4\|galm | 1.007144714127395E-12 |

| | | | | |
|---|---|---|---|---|
| 18 | Electron transport chain | WP:WP1339 | pop4\|sez6l2\|si:dkeyp-75h12.5\|imp3\|lmbrd2b\|psmc3\|ENSGACG00000009179\|ENSGACG00000001473\|gpx4b\|tmco1\|rpl24\|rpl28\|rpl29\|mrps24\|ENSGACG00000017620\|TMEM250\|fbl\|si:dkey-26c10.5\|fance\|lamtor4\|ENSGACG00000013492\|rpl7l1\|efhc1\|grnb\|ENSGACG00000010600\|clpp\|ogal\|rpl6\|tbc1d31\|gemin2 | 1.8668330343694776E-11 |
| 18 | Formation of the ternary complex, and subsequently, the 43S complex | REAC:R-DRE-72695 | eif2s3\|rangrf\|surf1\|ENSGACG00000007324\|gstr\|sgsm1a\|si:dkey-119f1.1\|FUCA1\|FAM135B\|rps28\|KCNMA1\|psma5\|msra\|rpl31\|mrps10\|psmd6\|ENSGACG00000009156\|cnot1\|si:dkey-266j7.2 | 9.977014071182463E-10 |
| 18 | ribosomal subunit | GO:0044391 | gstr\|FUCA1\|zbtb8a\|usp32\|march7\|ndufb10\|fynb\|ANO4\|galm | 1.2025780116575225E-9 |
| 18 | cytosolic large ribosomal subunit | GO:0022625 | zbtb8a\|usp32\|march7\|ndufb10\|fynb\|ANO4\|galm | 2.7927629137733214E-9 |
| 18 | Oxidative phosphorylation | WP:WP1335 | si:dkeyp-75h12.5\|ENSGACG00000009179\|ENSGACG00000001473\|gpx4b\|rpl24\|rpl29\|mrps24\|rpl11\|ENSGACG00000017620\|TMEM250\|nbeaa\|fbl\|fance\|lamtor4\|mrpl3\|ENSGACG00000013492\|rpl7l1\|efhc1\|clpp\|rpl6\|tbc1d31 | 3.0212567419117264E-9 |
| 18 | cytoplasm | GO:0005737 | hcfc1a\|gstr\|srp9\|FUCA1\|itsn2a\|ulk1b\|ENSGACG00000016474\|elmod2\|ENSGACG00000015390\|zbtb8a\|SNX14\|arl1\|ythdf3\|usp48\|usp47\|nme2a\|cntn3a.1\|dnajc15\|zdhhc17\|arnt\|hesx1\|ENSGACG00000012963\|gstk4\|usp32\|march7\|NDUFA4\|ndufb10\|fynb\|sub1b\|dhrs13a.3\|psmb7\|vps28\|ANO4\|mpv17\|ENSGACG00000002513\|rpl13\|vma21\|psmd9\|cyc1\|llph\|si:dkey-26c10.5\|uqcrq\|tpst1\|fkbp2\|e | 5.5687443372667445E-9 |

| | | | | |
|---|---|---|---|---|
| | | | if3k\|ENSGACG00000015644\|cyb5b\|eif1b\|galm\|ENSGACG00000019647\|ENSGACG00000015285\|ENSGACG00000018556\|chka\|ENSGACG00000004591 | |
| 18 | Ribosomal scanning and start codon recognition | REAC:R-DRE-72702 | eif2s3\|rangrf\|surf1\|ENSGACG00000007324\|gstr\|sgsm1a\|si:dkey-119f1.1\|FUCA1\|FAM135B\|rps28\|KCNMA1\|psma5\|msra\|rpl31\|mrps10\|psmd6\|ENSGACG00000009156\|cnot1\|si:dkey-266j7.2 | 6.2127749711676025E-9 |
| 18 | large ribosomal subunit | GO:0015934 | zbtb8a\|usp32\|march7\|ndufb10\|fynb\|ANO4\|galm | 2.4712823439946277E-7 |
| 18 | cytosol | GO:0005829 | hcfc1a\|gstr\|srp9\|FUCA1\|itsn2a\|zbtb8a\|ythdf3\|nme2a\|usp32\|march7\|ndufb10\|fynb\|psmb7\|ANO4\|vma21\|tpst1\|galm\|ENSGACG00000015285 | 2.8302335216694976E-7 |
| 18 | Spliceosome | KEGG:03040 | nubp1\|ENSGACG00000015196\|cx40.8\|ap3d1\|tmem63c\|ENSGACG00000008852\|arl2\|si:ch211-51h9.6\|ccdc90b\|mmgt1\|papolg\|smdt1b\|mettl5\|adcy3a\|ENSGACG00000002373\|PPFIA3\|tegt\|rnf13\|chchd7\|eif3g\|rpl3\|si:dkey-250d21.1\|sqstm1\|pabpc1b\|aimp1\|tmem50a | 9.035999388890336E-7 |
| 18 | translation | GO:0006412 | gstr\|FUCA1\|FAM135B\|zbtb8a\|gstk4\|usp32\|march7\|NDUFA4\|ndufb10\|fynb\|ANO4\|ENSGACG00000002513\|galm | 1.128091918646213E-6 |
| 18 | Metabolism of proteins | REAC:R-DRE-392499 | eif2s3\|top2b\|plxna4\|nudt5\|rangrf\|surf1\|ENSGACG00000007324\|gstr\|ENSGACG00000004293\|sgsm1a\|si:dkey-119f1.1\|FUCA1\|FAM135B\|rps28\|rps24\|SUMF2\|KCNMA1\|h2afva\|GRIA3\|etfb\|sec24c\|sp3b\|gnb5b\|ENSGACG00000007357\|si:ch211-14a17.11\|smim19\|NDUFB1\|smim15\|chmp3\|gstk4\|imp3 | 3.555718316687005E-6 |

| | | | | |
|---|---|---|---|---|
| | | | \|usp32\|march7\|emc10\|gm2a\|ndufb10\|micu3a\|usp20\|wdr60\|fam32a\|ano11\|psmb3\|psma3\|psma5\|msra\|ENSGACG00000016901\|tiprl\|wdr18\|tecpr2\|rpl39\|rpl30\|rpl34\|rpl31\|suclg1\|PDCL3\|mrps16\|mrps10\|psmd6\|ENSGACG00000009156\|arl6ip4\|cnot1\|si:dkey-266j7.2\|eif5b\|grk4\|ENSGACG00000015228\|ENSGACG00000014564\|C8orf82\|ENSGACG00000011934\|c18h3orf33\|eef1b2\|nol12\|sh3gl2a | |
| 18 | Aerobic respiration and respiratory electron transport | REAC:R-DRE-1428517 | cbr1l\|imp3\|naa50\|lmbrd2b\|psmc3\|ENSGACG00000001473\|tmco1\|rpl24\|rpl28\|rpl29\|mrps24\|ENSGACG00000017620\|TMEM250\|nbeaa\|lamtor4\|snrnp70\|mrpl3\|rpl7l1\|grnb\|rpl6\|tbc1d31\|gemin2 | 3.058945064361065E-5 |
| 18 | Respiratory electron transport | REAC:R-DRE-611105 | cbr1l\|imp3\|naa50\|lmbrd2b\|psmc3\|tmco1\|rpl24\|rpl28\|rpl29\|mrps24\|lamtor4\|snrnp70\|mrpl3\|rpl7l1\|rpl6\|tbc1d31 | 6.694077275015651E-4 |
| 18 | multicellular organism development | GO:0007275 | mrpl28\|FUCA1\|itsn2a\|ift172\|bdh2\|ulk1b\|elmod2\|atpaf1\|ENSGACG00000019377\|zdhhc17\|usp32\|ndufb10\|prep\|sub1b\|pex11g\|ANO4\|mrps30\|rpl13\|rpl14\|llph\|eif3k\|eif4h\|ENSGACG00000016379\|mtap\|ENSGACG00000015285\|rpl7\|rap1ab\|nol12\|ENSGACG00000004591 | 0.002512 |
| 18 | multicellular organismal process | GO:0032501 | mrpl28\|FUCA1\|itsn2a\|ift172\|bdh2\|ulk1b\|elmod2\|atpaf1\|ENSGACG00000019377\|zdhhc17\|usp32\|NDUFA4\|ndufb10\|prep\|sub1b\|aqr\|pex11g\|ANO4\|mrps30\|sptbn1\|rpl13\|rpl14\|llph\|eif3k\|eif4h\|ENSGACG00000016379\|mtap\|ENSGACG00000015285\|rpl7\|rap1ab\|nol12\|ENSGACG00000004591 | 0.009548 |
| 18 | cell morphogenesis | GO:0000902 | mrpl28\|itsn2a\|ift172\|ulk1b\|elmod2\|atpaf1\|pex11g\|mrps30\|rpl14\|ENSGACG000000 | 0.012724716977182936 |

| | | | | |
|---|---|---|---|---|
| | | | 16379\|ENSGACG00000015285 | |
| 18 | developmental process | GO:0032502 | mrpl28\|FUCA1\|itsn2a\|ift172\|bdh2\|ulk1b\|elmod2\|atpaf1\|zbtb8a\|ENSGACG00000019377\|nme2a\|zdhhc17\|usp32\|NDUFA4\|ndufb10\|prep\|sub1b\|pex11g\|ANO4\|mrps30\|rpl13\|rpl14\|vma21\|llph\|eif3k\|eif4h\|ENSGACG00000016379\|mtap\|ENSGACG00000015285\|rpl7\|rap1ab\|nsa2\|nol12\|ENSGACG00000004591 | 0.012833501792404937 |
| 18 | Cristae formation | REAC:R-DRE-8949613 | ENSGACG00000001473\|ENSGACG00000017620\|TMEM250\|nbeaa\|grnb\|gemin2 | 0.015138620508903347 |
| 18 | Formation of ATP by chemiosmotic coupling | REAC:R-DRE-163210 | ENSGACG00000001473\|ENSGACG00000017620\|TMEM250\|nbeaa\|grnb\|gemin2 | 0.015138620508903347 |
| 18 | anatomical structure development | GO:0048856 | mrpl28\|FUCA1\|itsn2a\|ift172\|bdh2\|ulk1b\|elmod2\|atpaf1\|zbtb8a\|ENSGACG00000019377\|nme2a\|zdhhc17\|usp32\|NDUFA4\|ndufb10\|prep\|sub1b\|pex11g\|ANO4\|mrps30\|rpl13\|rpl14\|vma21\|llph\|eif3k\|eif4h\|ENSGACG00000016379\|mtap\|ENSGACG00000015285\|rpl7\|rap1ab\|nol12\|ENSGACG00000004591 | 0.015220197684692625 |
| 18 | Mitochondrial biogenesis | REAC:R-DRE-1592230 | ctdsplb\|ENSGACG00000001473\|ENSGACG00000017620\|TMEM250\|nbeaa\|grnb\|gemin2 | 0.015251717779146811 |
| 18 | anatomical structure morphogenesis | GO:0009653 | mrpl28\|itsn2a\|ift172\|bdh2\|ulk1b\|elmod2\|atpaf1\|ENSGACG00000019377\|zdhhc17\|NDUFA4\|prep\|pex11g\|mrps30\|rpl14\|llph\|eif4h\|ENSGACG00000016379\|ENSGACG00000015285\|rpl7\|nol12 | 0.023064937805495896 |
| 18 | catalytic activity | GO:0003824 | sgip1b\|srp9\|itsn2a\|ENSGACG00000019975\|rps16\|bdh2\|ulk1b\|ENSGACG00000015390\|SNX14\|abhd14b\|arl1\|eef1g\|ythdf3\|usp48\|dnajc15\|hesx1\|psmb7\|aqr\|vps28\|uap1l1\|pex11g\|mrps30\|prkn\|mrps27\|rpl13\|vma21\|llph\|eif | 0.030376302871426293 |

| | | | | |
|---|---|---|---|---|
| | | | 3k\|cyb5b\|ENSGACG00000012151\|ENSGACG00000019647\|mtap\|ENSGACG00000018556\|rap1ab\|nfasca | |
| 18 | cytosolic small ribosomal subunit | GO:0022627 | gstr\|FUCA1 | 0.036530223300403036 |
| 18 | demethylase activity | GO:0032451 | rps16\|bdh2\|mrps30 | 0.038450575912334556 |
| 18 | Proteasome | KEGG:03050 | mrps18a\|phb\|sntg1\|adarb1b\|sec24c\|ddx47\|aar2\|picalma\|mtf1\|ENSGACG00000003280 | 0.039927 |
| 18 | embryo development | GO:0009790 | mrpl28\|FUCA1\|elmod2\|ENSGACG00000019377\|usp32\|ndufb10\|mrps30\|llph\|eif4h\|rpl7\|nol12 | 0.046500934221130524 |
| Overlap | Electron transport chain | WP:WP1339 | sgip1b\|eci1\|ndufa10\|cbr1l\|smc3\|ENSGACG00000015512\|atp5f1d\|eif4g3a\|lnpep\|pex11g\|syncrip\|sf3b4\|mrpl44 | 6.85189818650875E-10 |
| Overlap | Oxidative phosphorylation | KEGG:00190 | atp6v1f\|sgip1b\|eci1\|cbr1l\|smc3\|ENSGACG00000015512\|vps8\|atp5f1d\|eif4g3a\|lnpep\|ENSGACG00000015371\|pex11g\|ptrhd1\|syncrip\|sf3b4\|mrpl44 | 3.049936054079204E-9 |
| Overlap | Oxidative phosphorylation | WP:WP1335 | sgip1b\|eci1\|ENSGACG00000015512\|vps8\|eif4g3a\|lnpep\|pex11g\|syncrip\|mrpl44 | 3.7990165548555965E-7 |
| Overlap | Spliceosome | KEGG:03040 | ptgesl\|ACSL1\|aqr\|vps16\|ENSGACG00000002175\|bdh2\|mgst3b\|pgap1\|kifap3a\|fth1b | 0.001791 |
| Overlap | cytoplasm | GO:0005737 | chia.1\|afap1\|nupr1a\|hivep1\|gstr\|farsa\|zgc:163098\|atp5f1c\|gpx4b\|FAM126A\|ENSGACG00000012206\|si:ch211-282b22.1\|lrp1ab\|mrpl35 | 0.003333 |
| Overlap | Formation of ATP by chemiosmotic coupling | REAC:R-DRE-163210 | cbr1l\|pex11g\|sf3b4 | 0.033399 |
| Overlap | Cristae formation | REAC:R-DRE-8949613 | cbr1l\|pex11g\|sf3b4 | 0.033399 |
| Overlap | cytoplasmic translation | GO:0002181 | farsa\|atp5f1c | 0.040228 |

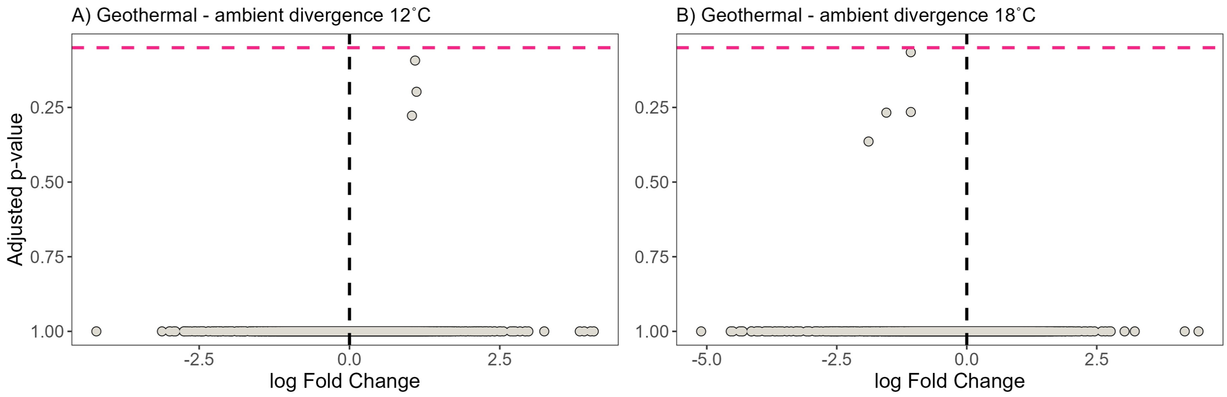


**Figure S1.** Gene expression divergence between geothermal and ambient ecotypes in the liver at A) 12°C and B) 18°C. No genes were differentially expressed in the liver.

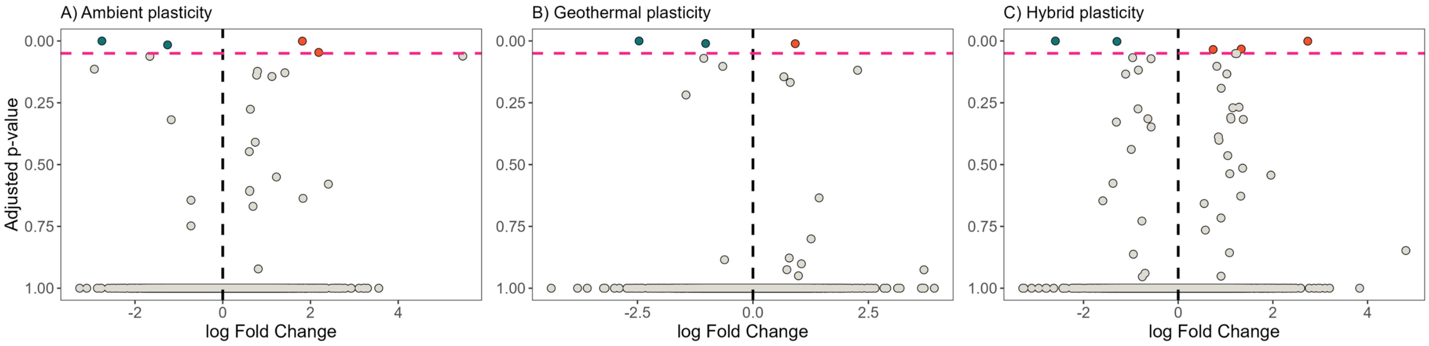


**Figure S2.** Gene expression plasticity between 12 and 18°C in the liver for the A) ambient ecotype, B) Geothermal ecotype, and C) hybrids. The pink line denotes significantly up- or down-regulated genes shown in red and blue, respectively.

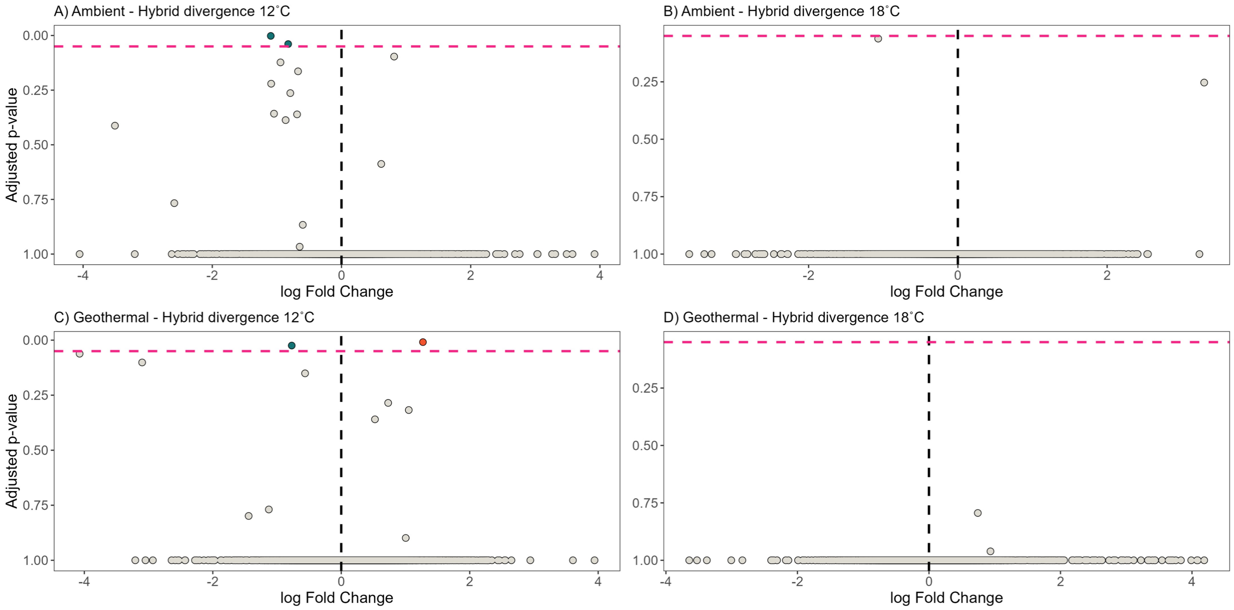


**Figure S3.** Divergence in gene expression between A) the ambient ecotype and hybrids at 12°C, B) the ambient ecotype and hybrids at 18°C, C) the geothermal ecotype and hybrids at 12°C, and D) the geothermal ecotype and hybrids at 18°C in the liver. The pink line denotes significantly up- or down-regulated genes shown in red and blue, respectively.

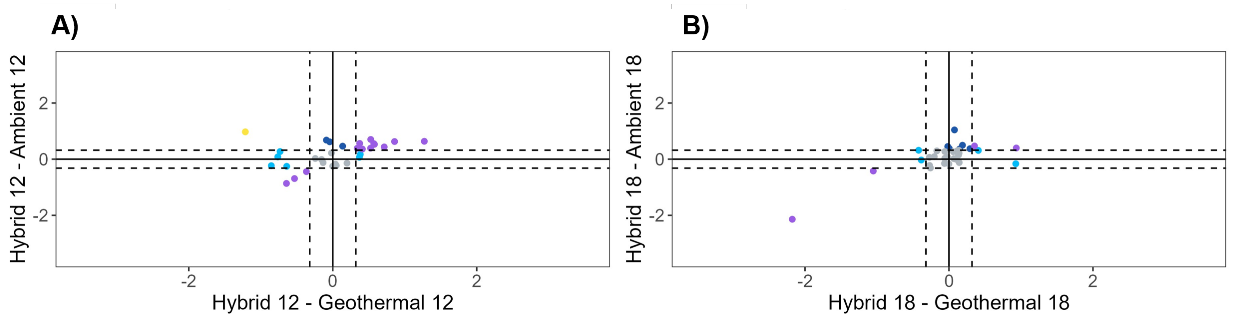


**Figure S4.** Inheritance patterns of divergent genes in hybrids compared to pure-strain geothermal and ambient ecotypes in the liver. Patterns of inheritance based on log fold change between hybrids relative to each pure-strained ecotype. Dominant inheritance (light and dark blue) patterns are between +/- 0.32 for each comparison. Additive inheritance (yellow) is defined as >0.32 in geothermal but <0.32 in the ambient ecotype or <0.32 in the geothermal ecotype but >0.32 in the ambient ecotype. Transgressive inheritance (purple) is defined as either < or > 0.32 in each of the pure-strained ecotypes. A) Patterns of inheritance in all differentially expressed genes at 12˚C. B) Patterns of inheritance in all differentially expressed genes at 18˚C.

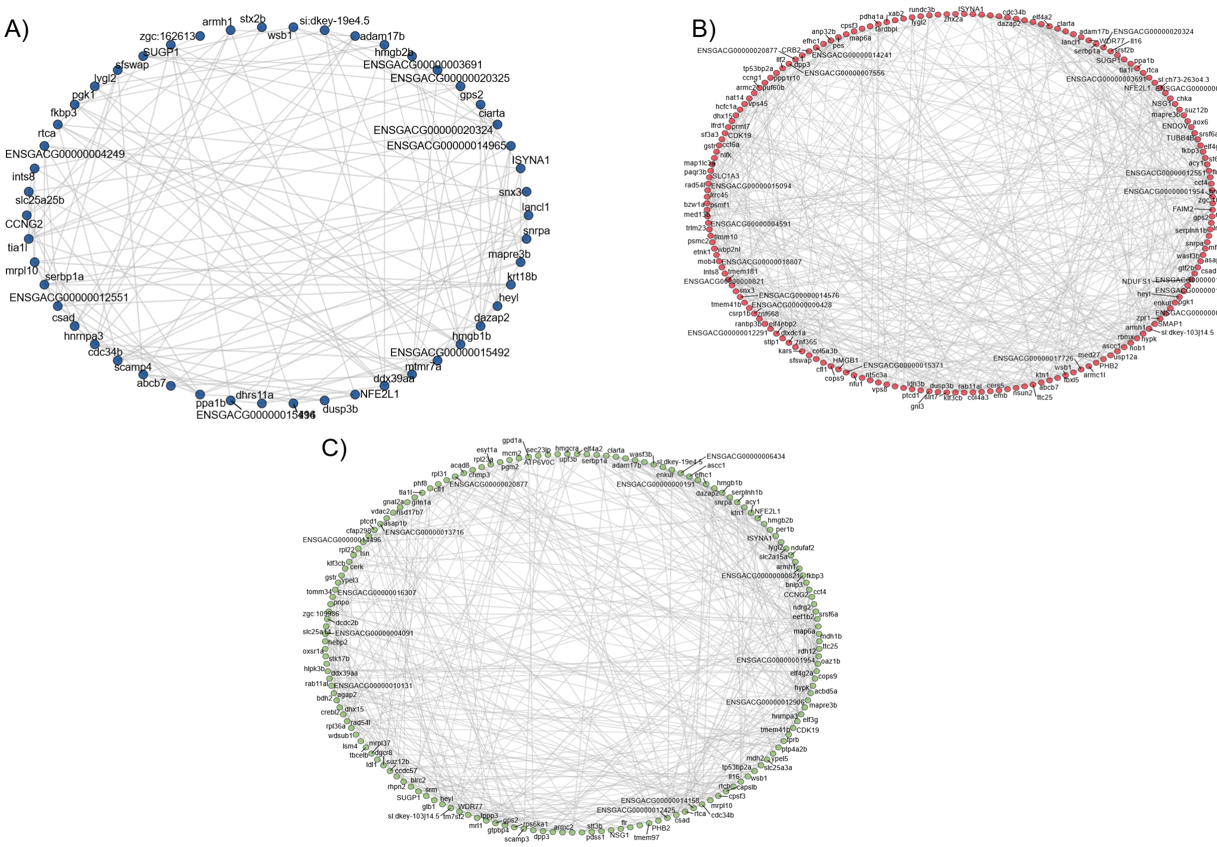


**Figure S5.** Whole gene networks associated with plastic responses to 12 and 18°C for the A) ambient ecotypes, B) geothermal ecotypes, and C) hybrids in the brain.

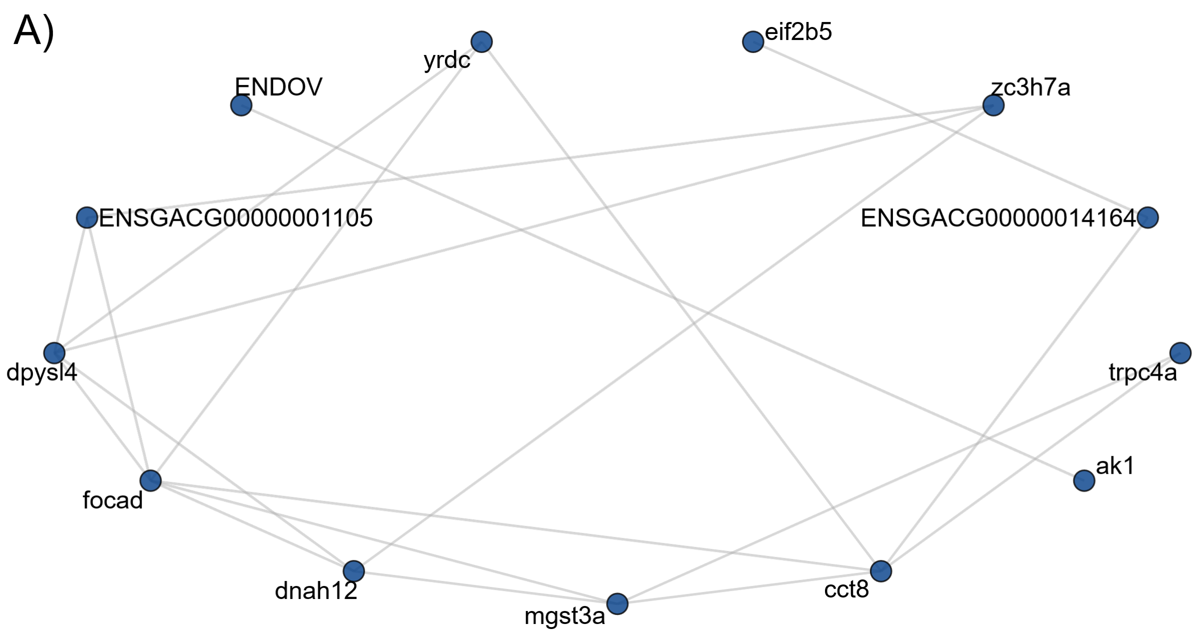


**Figure S6.** Gene network divergence between geothermal and ambient ecotypes at 12°C in the liver.

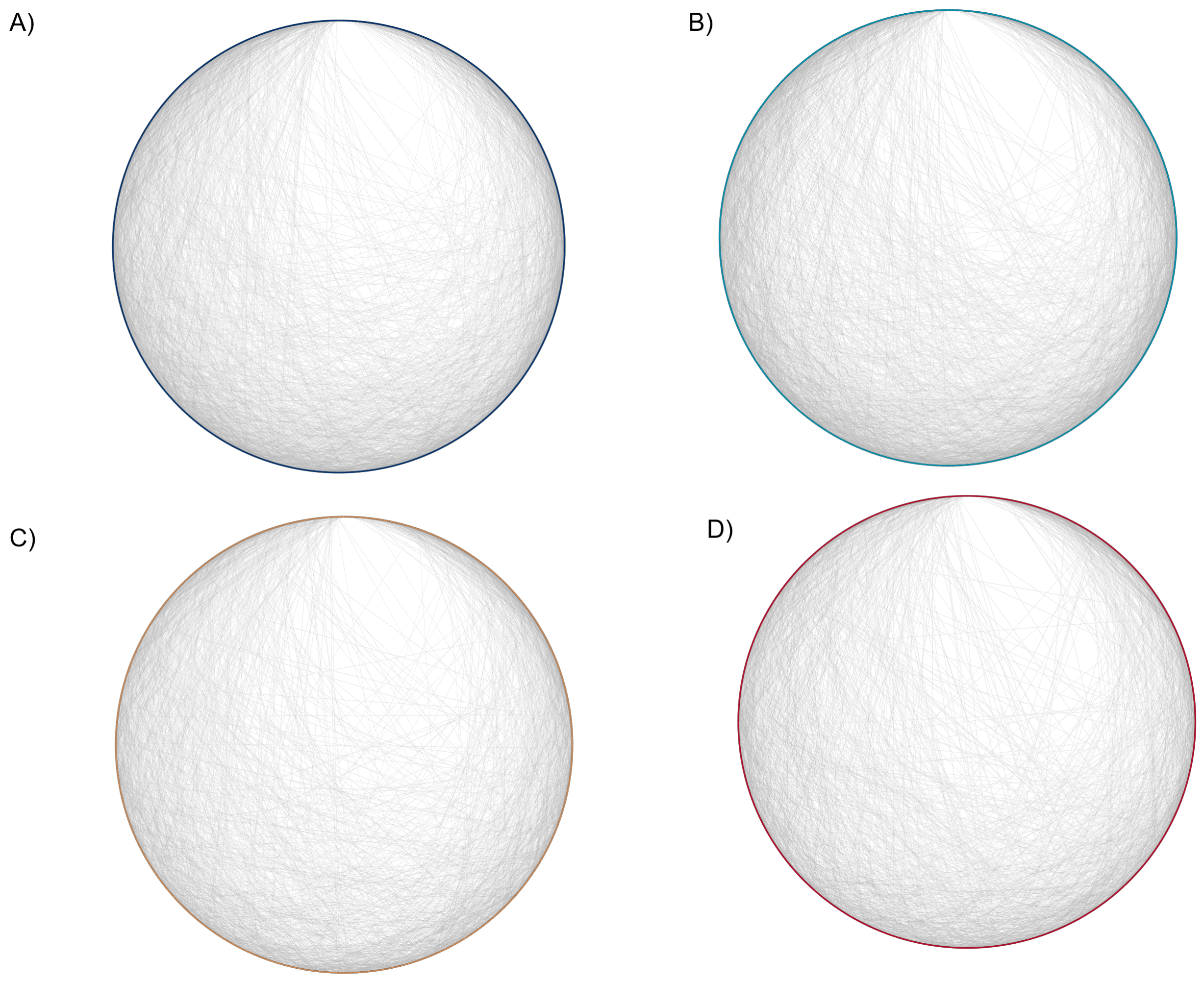

**Figure S7.** The gene networks associated with divergence between pure-strain ecotypes and hybrids A) ambient vs hybrid at 12°C, B) ambient vs hybrid at 18°C, C) geothermal vs hybrid at 12°C, and D) geothermal vs hybrid at 18°C in the brain.
